# Sensory neurons encode long-term inflammatory memory that promotes gastric regeneration and tumorigenesis

**DOI:** 10.64898/2026.06.15.732199

**Authors:** Yi Zeng, Puran Zhang, Feijing Wu, Ruhong Tu, Xiaofei Zhi, Hiroki Kobayashi, Jin Qian, Yosuke Ochiai, Biyun Zheng, Hualong Zheng, Shuang Li, Juli Lin, Masahiro Hata, Quin T. Waterbury, Junya Arai, Leah B. Zamechek, Timothy C. Wang

## Abstract

Inflammatory memory has emerged as a fundamental principle by which prior injury shapes future tissue responses, yet whether sensory neurons participate in long-term tissue memory remains unknown. Here, we show that vagal sensory neurons acquire a durable, experience-dependent state following gastric injury or *Helicobacter pylori* infection, leading to enhanced regeneration, metaplasia, and tumor progression upon re-injury. This neuronal program is stable, functionally transferable, and sufficient to drive epithelial responses in vivo. Mechanistically, injury-activated ILC2s establish sensory neuronal memory through IL-13–dependent epigenetic remodeling, inducing SMYD4-mediated H3K4 trimethylation and promoting CGRP-dependent activation of gastric epithelial cells. Together, our findings support a model in which tissue memory is not restricted to epithelial or immune compartments but emerges through coordinated long-term adaptations across multiple cellular systems. Within this framework, sensory neurons provide a persistent substrate for recall responses, linking prior inflammatory experience to sustained epithelial plasticity and cancer susceptibility.

**HIGHLIGHTS:**

- Sensory neurons function as a durable compartment of tissue memory.
- CGRP–RAMP1 signaling couples neuronal memory to gastric stem cells.
- ILC2-derived IL-13 establishes sensory neuronal memory programs.
- SMYD4-mediated H3K4me3 stabilizes long-term neuronal memory.
- Neuronal memory promotes gastric regeneration and tumor susceptibility.

## INTRODUCTION

Inflammation-associated tissue adaptation is increasingly recognized as a long-term biological process rather than a transient response to injury^1,2^. In gastric cancer, eradication of *Helicobacter pylori (H. pylori*) substantially reduces inflammation and lowers cancer incidence^3,4^; however, elevated cancer risk persists for decades after bacterial clearance^5^. These observations suggest that tissues retain a long-lasting imprint of prior inflammatory injury that continues to shape future epithelial behavior despite apparent recovery.

Recent studies have established inflammatory memory as a fundamental principle in tissue biology^6,7^. In epithelial systems, stem and progenitor cells can acquire persistent epigenetic states that enhance regenerative responses upon recurrent injury^8–11^. Likewise, innate immune populations can adopt trained or adaptive inflammatory states following prior stimulation^11^. More recently, persistent inflammatory programs have also been described in innate lymphoid populations, including ILC2s^12^. Together, these findings suggest that tissue memory is not restricted to a single cellular compartment but instead may emerge through coordinated adaptations across epithelial and immune systems.

Although epithelial injury memory has been proposed to underlie enhanced tissue regeneration^8,9^, whether epithelial responses are fully cell-intrinsic remains incompletely understood. Tissue repair and regeneration occur within complex multicellular environments in which immune, stromal, and neural signals continuously shape epithelial behavior^7^. Thus, an important unresolved question is whether additional long-lived cellular systems contribute to the persistence, integration, or recall of prior inflammatory experience. Sensory neurons are uniquely positioned to fulfill such functions^13,14^. These cells continuously monitor tissue integrity, relay inflammatory information to the central nervous system, and persist for the lifetime of the organism^15^. Previous studies have demonstrated that sensory neurons remain activated following gastric inflammation and that inflammatory signals propagate from the stomach to the brainstem^16^. Unlike rapidly turning-over epithelial cells, sensory neurons possess anatomical stability and longevity necessary to potentially retain prior inflammatory experience over extended periods.

Here, we demonstrate that sensory neurons encode a durable inflammatory memory state that regulates long-term epithelial responses in the stomach. We show that this neuronal program is established through immune–neural interactions, maintained through IL-13–SMYD4–dependent epigenetic remodeling, and recalled upon recurrent tissue injury to promote regeneration and tumor progression. Our findings support a broader framework in which tissue memory is distributed across multiple cellular compartments, while sensory neurons provide a uniquely persistent and functionally transferable component of long-term tissue adaptation.

## RESULTS

### Prior gastric inflammation establishes durable inflammatory memory and accelerates regeneration and tumor progression

We studied prior gastric inflammation using a series of complementary injury and tumor models, including a high-dose tamoxifen (HDT)–induced gastric injury model^17^, *Helicobacter pylori* infection plus antibiotic eradication^4^, an acetic acid–induced gastric ulcer healing model^18^, and multiple gastric cancer models (including orthotopic^19^ and spontaneous^20,21^). In the HDT injury model, high-dose tamoxifen induced rapid loss of secretory (parietal and chief) cells accompanied by pyloric metaplasia/Spasmolytic Polypeptide–Expressing Metaplasia (SPEM) **(Fig 1A)**. Histological analysis confirmed that gastric cellular composition returned to baseline within 3 weeks, with complete tissue repair **(Fig S1A)**. Importantly, when HDT injury was re-administered after full recovery, these mice exhibited an accelerated regenerative response compared with mice experiencing injury for the first time **(Fig S1A)**. Quantitative analyses showed that in days 3 and 7 following re-injury, epithelial proliferation and metaplastic marker expression were significantly higher in recall mice than in controls. This recall response was characterized by more rapid expansion of ATP4B⁺ parietal cell and GIF⁺ chief cell lineages, earlier re-emergence of GSII⁺ GIF⁺ SPEM markers^22^, and enhanced Ki67⁺ glandular proliferation **(Fig 1B-1E)**. Staining CD45+ immune cells revealed a similar robust recruitment of infiltrating immune cells to the glandular isthmus during both injury and recall phases **(Fig S1B)**. This enhanced responsiveness was durable, such that even when the second injury was administered six months after the initial insult, mice retained a robust recall phenotype, consistent with a long-lasting memory state within the gastric tissue **(Fig S1D and S1E)**.

**Figure 1.**
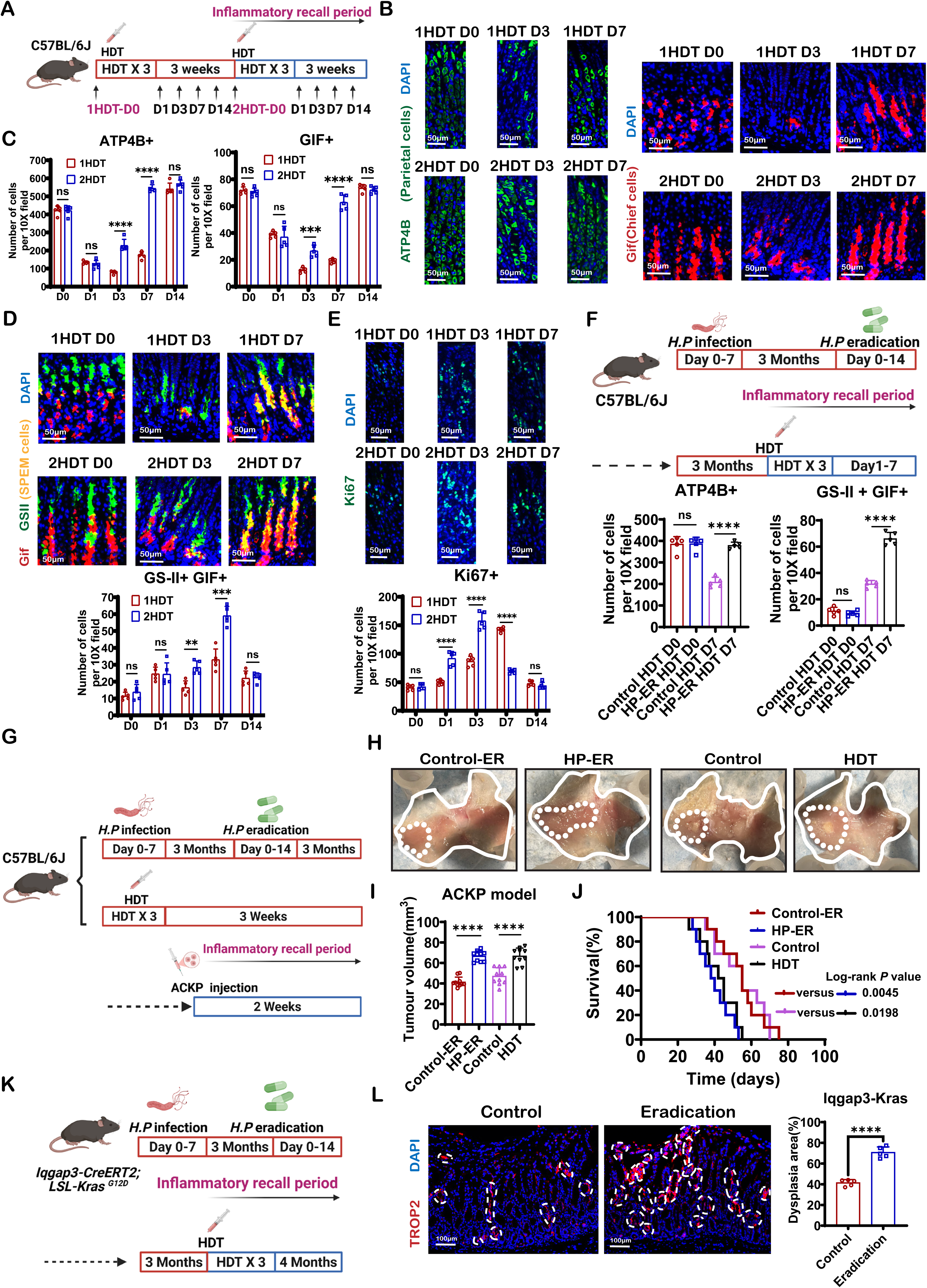
Prior gastric inflammation establishes durable inflammatory memory that accelerates regeneration and tumor progression. **(A)** Schematic of the HDT injury and recall model. **(B)** Representative immunofluorescence images of ATP4B and GIF staining in gastric glands after 1HDT or 2HDT. DAPI marks nuclei. Scale bars, 50 µm. **(C)** Quantification of ATP4B^+^ parietal cells, GIF^+^ chief cells and GSII^+^GIF^+^ SPEM cells per ×10 field after 1HDT and 2HDT (n = 5 mice per group). **(D)** Representative GSII and GIF immunofluorescence staining with quantification of GSII^+^GIF^+^ SPEM cells per ×10 field after 1HDT or 2HDT (n = 5 mice per group). Scale bars, 50 µm. **(E)** Representative Ki67 immunofluorescence staining with quantification of epithelial cells after 1HDT or 2HDT (n = 5 mice per group). Scale bars, 50 µm. **(F)** Schematic of *H. pylori* infection, antibiotic eradication and HDT challenge, with quantification of ATP4B^+^ and GSII^+^GIF^+^ cells (n = 5 mice per group). HP, *H. pylori*; ER,eradication. **(G)** Schematic for assessing tumor progression after *H. pylori*- or HDT-induced inflammatory memory. **(H)** Representative gross images of orthotopic ACKP tumors from the indicated groups. **(I)** Quantification of tumor volume in the orthotopic ACKP model (n = 10 mice per group). **(J)** Kaplan–Meier survival curves for the indicated groups (n = 10 mice per group). **(K)** Schematic for evaluating dysplasia in *Iqgap3-CreERT2; LSL-Kras^G12D^* mice. **(L)** Representative TROP2 immunofluorescence staining with quantification of dysplasia area (%) (n = 5 mice per group). Scale bars, 100 µm. Statistical significance indicated by \**p* < 0.05, \*\**p* < 0.01, \*\*\**p* < 0.001, \*\*\*\**p* < 0.0001; ns, not significant. Data are mean ± s.e.m. *P* values were calculated using two-sided t-tests (Figure 1C**,1D, 1E,1F,1I and 1L)** and log-rank test (Figure 1J). See also **Figure S1** and **S2**.

In a clinically relevant model of *H. pylori* infection, infected mice with chronic gastritis were successfully treated with antibiotic eradication with restoration of normal gastric histology **(Fig S1F and S1G)**, HDT challenge elicited accelerated regeneration and metaplasia in mice post-eradication, resembling the recall phenotype observed after prior HDT injury **(Fig 1F and S1H)** and demonstrating that inflammatory memory can persist after pathogen clearance and gastritis resolution. We also employed an acetic acid–induced gastric ulcer healing model in mice with a history of HDT injury or *H. pylori* eradication, and found significantly faster ulcer healing, with nearly complete healing of ulcers by day 7, consistent with inflammatory memory **(Fig S1I and S1J)**.

However, inflammatory memory imposed a clear cost in cancer susceptibility. Across multiple gastric cancer models, including orthotopic implantation of ACKP tumors **(Fig 1G-1)**, spontaneous intestinal-type gastric cancer^20^ **(Fig 1K and 1L)**, and diffuse-type gastric cancer models^21^ **(Fig S2A-S2C)**, mice with a history of prior gastric inflammation exhibited markedly accelerated tumor progression and increased tumor burden compared with control mice. Notably, this enhanced tumor susceptibility translated into a significant reduction in overall survival (**Fig 1J and S2E**). Together, these results demonstrate that prior gastric inflammation establishes a durable memory state that accelerates tissue regeneration upon re-injury while simultaneously creating a permissive environment for tumor development.

To determine whether epithelial cells alone retain an intrinsic inflammatory memory state following *H. pylori* exposure, we isolated Iqgap3⁺ gastric epithelial cells from mice with or without prior inflammatory experience and established organoid cultures **(Fig S2E)**. Although organoids derived from previously infected mice exhibited a modest trend toward increased growth at later time points, no significant differences in organoid size were observed compared with inflammation-naive controls **(Fig S2F)**. These findings suggest that the durable inflammatory memory phenotype observed in vivo cannot be fully explained by epithelial-intrinsic memory alone and likely requires additional support from the surrounding tissue microenvironment.

### Sensory neuron activation is associated with gastric inflammatory memory

Given that the stomach is a densely innervated organ with extensive peripheral nerve networks, we next investigated whether gastric inflammatory memory is associated with altered neural innervation during recall responses. To address this, we compared neural remodeling during the acute inflammatory phase (day 3 after HDT) between primary and secondary injury conditions. Using calcitonin gene-related peptide (CGRP) as a marker of peptidergic sensory neurons^23,24^, we found that CGRP⁺ sensory innervation was markedly increased during secondary inflammation compared with primary injury in both the HDT model and the *H. pylori* infection–eradication model **(Fig 2A and S3A)**. In contrast, adrenergic (TH+) and cholinergic (ChAT+) innervation remained unchanged **(Fig S3B)**. Moreover, immunostaining for another major peptidergic neuropeptide, Substance P, revealed no significant differences between primary and secondary inflammatory responses **(Fig S3C)**, suggesting a selective remodeling of CGRP⁺ sensory neural circuits during gastric inflammatory memory. This phenomenon was consistently observed across regeneration and tumor models and was most pronounced in regions exhibiting accelerated regeneration and metaplasia **(Fig S3D-S3G)**, suggesting that prior inflammation primes sensory innervation for exaggerated responses upon re-injury.

**Figure 2.**
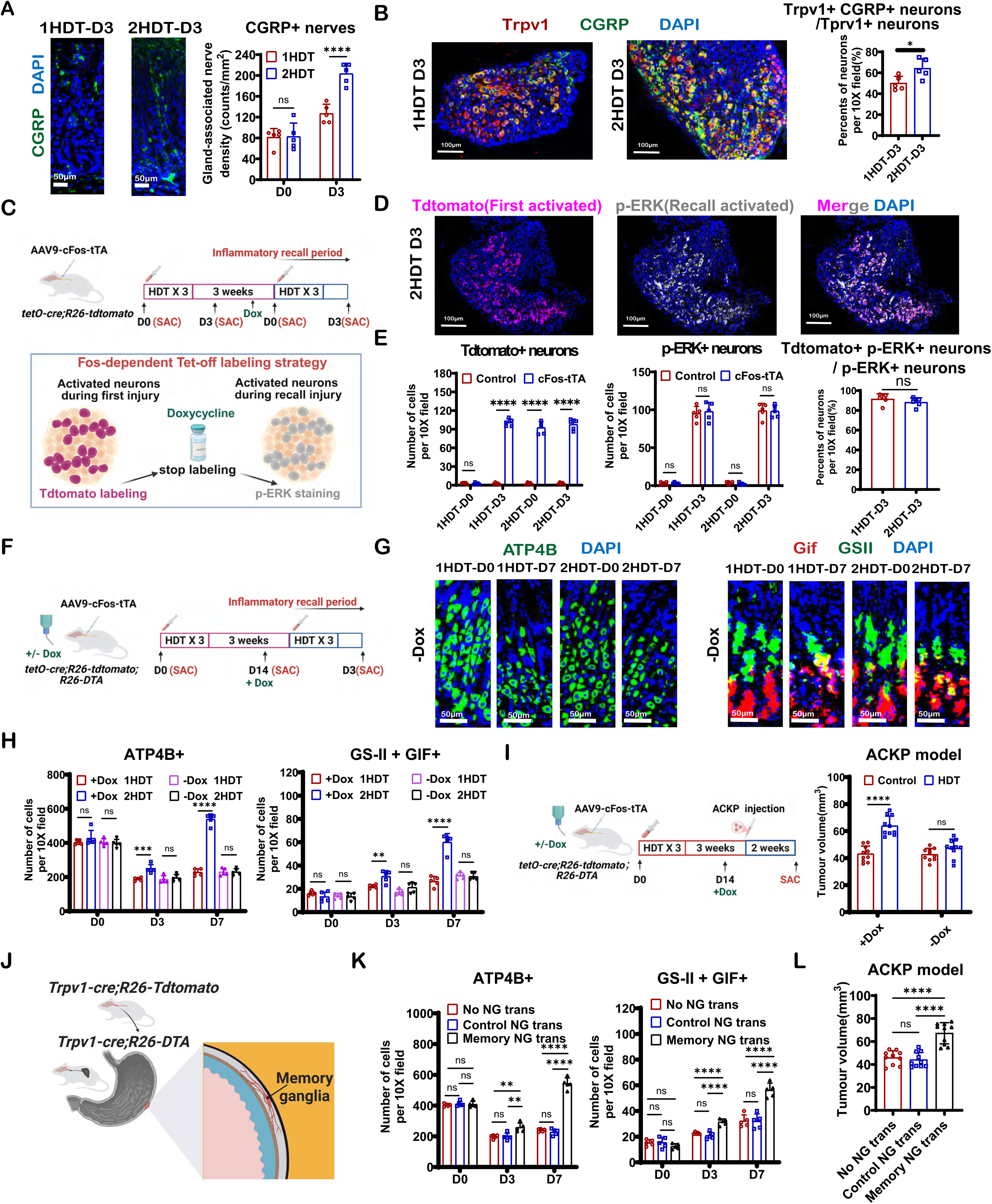
Sensory neuron activation is associated with gastric inflammatory memory. **(A)** Representative immunofluorescence images and quantification of CGRP+ nerve density in gastric lesions after primary injury (1HDT) or recall injury (2HDT) (n = 5 mice per group). Scale bars, 50 µm. **(B)** Representative immunofluorescence images of nodose ganglia stained for Trpv1 and CGRP with quantification after 1HDT or 2HDT (n = 5 mice per group). Scale bars, 100 µm. **(C)** Schematic of the cFos-dependent Tet-off strategy used to label neurons activated during the first injury and identify reactivated neurons during inflammatory recall. **(D)** Fos-dependent Tet-off tracing in NG. Neurons activated during the initial injury are labeled by tdTomato, whereas neurons activated upon re-injury are marked by p-ERK. Representative immunofluorescence images show tdTomato^+^/p-ERK^+^ overlap. Scale bars, 100 µm. **(E)** Quantification of tdTomato^+^ and p-ERK^+^ neurons in Fos-dependent Tet-off traced NG. Scale bars, 100 µm (n =5 mice per group). **(F)** Experimental schematic of Fos-dependent targeting in *tetO-Cre; R26-tdTomato; R26-DTA* mice under doxycycline-on or -off conditions, followed by HDT injury. **(G, H)** Representative immunofluorescence images and quantification of ATP4B and GIF/GSII staining of gastric glands in *tetO-Cre; R26-tdTomato; R26-DTA* mice following Fos-dependent targeting of neurons activated during the first HDT injury (1HDT) and subsequent recall injury (2HDT) under doxycycline-on or -off conditions (n =5 mice per group). Scale bars, 50 μm. **(I)** Experimental schematic of Fos-dependent targeting in *tetO-Cre; R26-tdTomato; R26-DTA* mice under doxycycline-on or -off conditions, followed by orthotopic ACKP tumor implantation, with tumor volume quantified (n = 10 mice per group). **(J)** Schematic of nodose ganglion transplantation from inflammation-experienced or control donors into *Trpv1-Cre; R26-DTA* recipients. **(K)** Quantification of ATP4B^+^ cells and GSII^+^GIF^+^ cells in gastric glands with or without surrounding transplanted ganglia (n = 5 mice per group). **(L)** Quantification of tumor burden in mice with or without surrounding transplanted ganglia (n = 10 mice per group). Statistical significance indicated by \**p* < 0.05, \*\**p* < 0.01, \*\*\**p* < 0.001, \*\*\*\**p* < 0.0001; ns, not significant. Data are mean ± s.e.m. *P* values were calculated using one-way ANOVA with Dunnett’s test (Figure 2K **and 2L)** and two-sided t-tests (Figure 2B**, 2E, 2H and 2I)**. See also **Figure S3** and **S4**.

At the level of sensory ganglia, c-Fos staining revealed a significant increase in activated neurons in the nodose ganglia (NG) or Jugular–Nodose Complex (JNC) following recall injury, whereas dorsal root ganglia (DRG) did not show a comparable increase **(Fig S3H)**. Consistently, CGRP expression in Trpv1⁺ NG neurons was elevated during recall **(Fig 2B and S3H-S3J)**, indicating that gastric inflammatory memory preferentially engages the vagal sensory pathway.

To directly test whether intact sensory innervation is required for inflammatory memory, we performed selective denervation. Unilateral transection of the anterior vagal trunk (UVT) ^25^caused localized loss of sensory innervation in the anterior gastric wall **(Fig S3K and S3L)**. In denervated regions, recall injury failed to induce accelerated regeneration, with delayed repair, reduced proliferation, and diminished metaplastic marker expression, whereas the innervated wall retained a robust recall phenotype **(Fig S3M)**. Similarly, denervation abolished memory-associated acceleration of tumor growth in tumor models **(Fig S3N)**, demonstrating that intact sensory innervation is required for the manifestation of gastric inflammatory memory and that this phenotype is spatially restricted to innervated regions.

To trace inflammatory memory to a specific neuronal population, we used a Fos-dependent Tet-off labeling strategy by injecting AAV-cFos-tTA^26^ into the NG of *tetO-Cre; R26-tdTomato*^27^ mice, thereby permanently labeling neurons activated during the initial injury. Doxycycline was subsequently administered to terminate further labeling **(Fig 2C)**. During recall injury, p-ERK staining revealed preferential reactivation of the tdTomato-labelled neuronal population, demonstrating substantial overlap between neurons activated during the initial and recall responses **(Fig 2D and 2E)**.

Finally, to determine whether enhanced peripheral activation is transmitted to central circuits, we examined neuronal activity in the nucleus tractus solitarius (NTS), the primary brainstem target of vagal afferents. Both c-Fos staining and Magnetic resonance imaging (MRI) analyses^28,29^ revealed significantly increased NTS activation during recall **(Fig S4A-S4C)**, indicating that peripheral inflammatory memory is conveyed to central sensory processing centers. Together, these results demonstrate that gastric inflammatory memory is associated with activation of a defined vagal sensory circuit, spanning enhanced peripheral innervation, selective NG activation, and propagation to central pathways.

### Trpv1⁺ memory-encoded sensory neurons are both necessary and sufficient for inflammatory memory recall

To determine whether activation of sensory neurons is sufficient to drive recall responses, we selectively activated Trpv1⁺ sensory neurons using a chemogenetic approach. Trpv1⁺ neurons were chosen because they constitute a major population of peptidergic visceral sensory neurons^23^ and are preferentially activated during inflammatory memory recall. Following gastric injury, chemogenetic activation of Trpv1⁺ neurons was sufficient to accelerate gastric epithelial regeneration, as evidenced by increased glandular proliferation, expansion of regenerative regions, and induction of metaplastic markers, closely phenocopying the recall response observed after prior inflammation **(Fig S4D)**.

To assess the necessity of Trpv1⁺ sensory neurons for inflammatory memory expression, we genetically ablated Trpv1⁺ neurons using *Trpv1-Cre; R26-DTA* mice, which eliminated the accelerated regenerative response during recall injury. In these mice, epithelial proliferation, metaplastic marker expression, and tissue repair kinetics following re-injury were comparable to those observed during primary injury in inflammation-naive controls, demonstrating that Trpv1⁺ sensory neurons are required for the expression of inflammatory memory **(Fig S4E)**.

To address whether inflammatory memory resides within a specific subset of neurons activated during the initial injury, we injected AAV-cFos-tTA into the NG of *tetO-Cre; R26-tdTomato; R26-DTA* mice to implement a Fos-dependent Tet-off–based activity-dependent neuronal ablation strategy. Neurons activated during the first HDT injury were permanently labeled and ablated, after which doxycycline was administered to terminate further labeling and ablation prior to recall **(Fig 2F)**. Selective ablation of these injury-activated neurons did not affect baseline gastric architecture or primary injury responses. However, during recall injury, mice lacking these memory-encoded neurons failed to exhibit accelerated regeneration or metaplasia; proliferative responses, regenerative marker expression, and repair kinetics were indistinguishable from those observed during primary injury **(Fig 2G and 2H)**. Similarly, in the HDT-preconditioned ACKP tumor model, ablation of neurons activated during the primary inflammatory episode abolished the enhanced tumor progression observed during recall, effectively eliminating the tumor-promoting memory phenotype **(Fig 2I)**.These results indicate that inflammatory memory is encoded within a small subset of sensory neurons activated during the initial inflammatory insult and that these neurons are required for recall responses.

To explore whether memory-encoded neurons were fully sufficient to transfer inflammatory memory, we next performed nodose ganglion transplantation experiments. Nodose ganglia from mice harboring inflammatory memory were transplanted ^30^ beneath the gastric serosa of recipient mice lacking Trpv1⁺ sensory neurons **(Fig 2J and S4H)**. Fluorescent labeling revealed that NGs with prior inflammatory experience exhibited enhanced axonal outgrowth after transplantation into the stomach, whereas graft survival rates were comparable between groups **(Fig S4I)**. Following HDT-induced glandular injury, memory-encoded NGs displayed sustained neuronal activation, as indicated by increased p-ERK signaling, and extended axonal projections toward the glandular epithelial isthmus region after transplantation into the stomach. In contrast, naive NGs exhibited markedly reduced axonal extension and epithelial targeting capacity under the same conditions **(Fig S4J)**. Transplantation of memory-encoded nodose ganglia markedly restored the accelerated regeneration and tumor-promoting effects in the gastric wall surrounding the graft site in recipient mice, whereas transplantation of inflammation-naive nodose ganglia failed to do so **(Fig 2K,2L and S4K, S4L)**. These findings demonstrate that inflammatory memory is stably maintained within a defined population of Trpv1⁺ sensory neurons and can be transferred across individuals independently of the original inflammatory environment. Together, these results establish that Trpv1⁺ sensory neurons are both necessary and sufficient for inflammatory memory recall and that inflammatory memory is encoded within a discrete ensemble of neurons activated during the initial inflammatory insult.

### CGRP–RAMP1 signaling converts sensory neuronal memory into epithelial regeneration and tumor growth

To elucidate how neuronal memory is translated into epithelial responses during recall, we focused on CGRP, a dominant sensory neuropeptide in the stomach and a candidate effector linking neuronal activation to epithelial behavior^14,31^. In immunodeficient *Rag2^⁻/⁻^γc^⁻/⁻^* mice lacking adaptive and innate lymphoid cells, administration of exogenous CGRP by intraperitoneal (i.p.) injection following HDT-induced injury markedly enhanced epithelial proliferation and regeneration, phenocopying recall responses observed after prior inflammation, suggesting that CGRP directly promotes epithelial regeneration independent of secondary inflammatory mechanisms **(Fig S5A-5C)**.

CGRP signals through a receptor complex containing RAMP1^32^. Immunostaining revealed that whereas RAMP1 was primarily localized to the gland base at baseline, it became markedly enriched in the gastric isthmus following HDT injury **(Fig 3A and 3B)**, the main site for gastric stem and progenitor cells. Consistently, RAMP1 expression was prominent in Iqgap3⁺ progenitor cells^20^ **(Fig S5D)**. Lineage tracing further showed that Iqgap3⁺ progenitors exhibited enhanced proliferative capacity and disproportionately contributed to gland regeneration during recall **(Fig 3C)**. Consistently, exogenous CGRP administration failed to accelerate glandular regeneration or metaplastic responses in mice with selective deletion of RAMP1 in gastric progenitors **(Fig S5E)**. Moreover, selective deletion of RAMP1 in gastric progenitors abolished inflammation memory–associated acceleration of regeneration without affecting baseline architecture or primary injury responses **(Fig 3D and S5F)**, indicating that CGRP–RAMP1 signaling directly regulates progenitor fate^31,33^ during recall.

**Figure 3.**
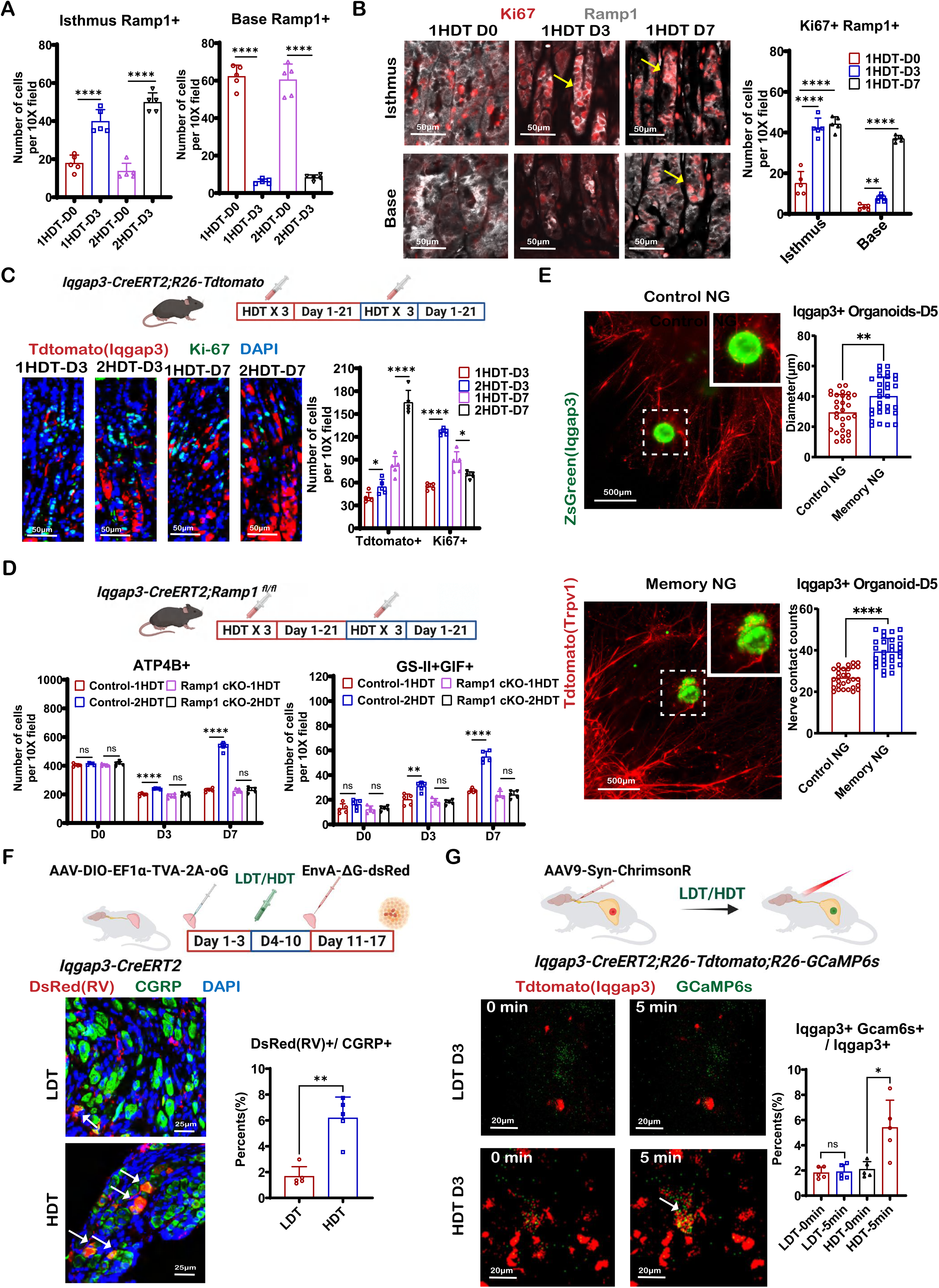
Sensory neuron–encoded inflammatory memory preferentially targets Ramp1^+^ gastric progenitor cells. **(A)** Quantification of Ramp1^+^ cells in gastric isthmus and base regions after primary injury (1HDT) or recall injury (2HDT) (n = 5 mice per group). **(B)** Representative immunofluorescence images of gastric glands stained for Ki67 and Ramp1 at D0, D3 and D7 after 1HDT with quantification of Ki67^+^Ramp1^+^ cells (n = 5 mice per group). Scale bars, 50 µm. **(C)** Schematic of Iqgap3+ cells lineage tracing and representative immunofluorescence images showing tdTomato-labelled Iqgap3-lineage cells with Ki67 staining at D3 and D7 after 1HDT or 2HDT with quantification of tdTomato^+^ and Ki67^+^ cells (n = 5 mice per group). Scale bars, 50 µm. **(D)** Schematic and quantification of ATP4B^+^ cells and GSII^+^GIF^+^ cells after 1HDT or 2HDT in *Iqgap3-CreERT2; Ramp1^flox/flox^* or *Iqgap3-CreERT2; Ramp1^+/+^* mice (n = 5 mice per group). **(E)** Gastric organoid–neuron co-culture showing tdTomato^+^ Trpv1-lineage neurons interacting with Iqgap3^+^ gastric organoids under control NG and memory NG with quantification of organoid diameter and nerve contacts (n = 30 spheroids/group). Scale bars, 500 µm. **(F)** Schematic of retrograde trans-synaptic tracing from gastric Iqgap3^+^ cells using EnvA-pseudotyped ΔG rabies virus showing labelled neurons in the nodose ganglion with high dose tamoxifen (HDT) or low dose tamoxifen (LDT). Representative immunofluorescence images of nodose ganglia stained for DsRed and CGRP with quantification after HDT or LDT (n = 5 mice per group). Scale bars, 25 µm. **(H)** Schematic of time-lapse Ca²⁺ imaging (GCaMP6s) and quantification in Iqgap3^+^ gastric cells following LDT or HDT stimulation. Scale bars, 20 µm. Statistical significance indicated by \**p* < 0.05, \*\**p* < 0.01, \*\*\**p* < 0.001, \*\*\*\**p* < 0.0001; ns, not significant. Data are mean ± s.e.m. *P* values were calculated using one-way ANOVA with Dunnett’s test (Figure 3B) or two-sided t-tests (Figure 3A**,3C,3D,3E,3F and 3G)**. See also **Figure S5** and **S6**.

Consistent with direct neuro–epithelial interactions, chemogenetic activation of Trpv1⁺ sensory neurons further potentiated gastric organoid growth and survival, whereas genetic deletion of RAMP1 in organoids completely abolished these effects, confirming a CGRP-dependent progenitor mechanism **(Fig S5G–S5I)**. Co-culture experiments further revealed that sensory neurons derived from memory-encoded NGs markedly enhanced the growth and survival of gastric organoids established from Iqgap3⁺ epithelial cells, whereas naive sensory neurons exhibited substantially weaker supportive effects **(Fig 3E)**. Notably, organoids co-cultured with memory-encoded NGs displayed increased direct contacts with sensory neurites, suggesting enhanced neuro–epithelial interactions during the recall state **(Fig 3E)**.

Whole-mount imaging and synaptophysin-based tracing^34^ identified CGRP⁺ sensory fibers terminating in the gastric isthmus adjacent to progenitor cells following HDT injury, suggesting the formation of specialized neuro–epithelial communication sites **(Fig S6A, S6B and Supplementary Video1)**. Blocking synaptic vesicles release from sensory neurons using tetanus toxin^35^ during the memory phase markedly attenuated recall-associated epithelial regeneration without altering the elevated neuronal activation or CGRP expression in the nodose ganglia **(Fig S6C-S6E)**, indicating that vesicle-mediated neurotransmission is required at the epithelial interface. Monosynaptic retrograde tracing was performed by selectively targeting Iqgap3⁺ gastric progenitors with EnvA-pseudotyped ΔG rabies virus^36^. Following HDT-induced injury, traced neurons were readily detected in the NG, confirming direct connectivity between gastric progenitors and vagal sensory neurons. In contrast, only rare traced neurons were observed following low-dose tamoxifen damage (LDT), indicating that robust sensory–epithelial circuit formation preferentially emerges under severe inflammatory injury conditions associated with inflammatory memory **(Fig 3F)**.

To determine whether neuronal activation can acutely modulate progenitor activity in vivo, we combined optogenetic activation^37^ of nodose ganglion neurons with real-time calcium imaging^38^ of Iqgap3⁺ progenitor cells. Following inflammatory injury, optogenetic stimulation elicited robust calcium influx in progenitor cells, whereas this response to neural stimulation was absent in mice without a prior inflammatory history **(Fig 3H and Supplementary Video2)**, supporting a model in which inflammation-conditioned sensory neurons establish rapid functional coupling with epithelial progenitors.

Consistent with these findings, administration of the CGRP receptor antagonist Rimegepant during the recall phase, after establishment of inflammatory memory, selectively attenuated memory-associated responses, including accelerated glandular regeneration, metaplastic progression, and enhanced tumor growth **(Fig. S6F–S6I)**. Notably, Rimegepant treatment also significantly improved survival in memory-bearing tumor models **(Fig S6J)**, highlighting CGRP–RAMP1 signaling as a critical effector pathway driving pathological recall responses during gastric inflammatory memory.

Together, these results demonstrate that CGRP–RAMP1–dependent neuro–epithelial communication is a key mechanism by which sensory neuronal memory is translated into epithelial regeneration and tumor growth.

### ILC2s instruct the establishment of sensory neuron–mediated inflammatory memory

To determine whether neuronal inflammatory memory depends on immune input, we examined its establishment in immunodeficient mice. In Rag1-deficient mice lacking adaptive immunity, CGRP⁺ sensory innervation and recall-associated regenerative and tumor-promoting responses were preserved **(Fig S7A-S7C)**, indicating that adaptive immunity is dispensable. In contrast, in *Rag2^⁻/⁻^γc^⁻/⁻^* mice lacking both adaptive and innate lymphoid cells, NG sensory neuron activation and CGRP⁺ innervation were markedly reduced, and the inflammatory memory phenotype was abolished **(Fig 4A, B and S7D)**, demonstrating a strict requirement for innate immune input.

**Figure 4.**
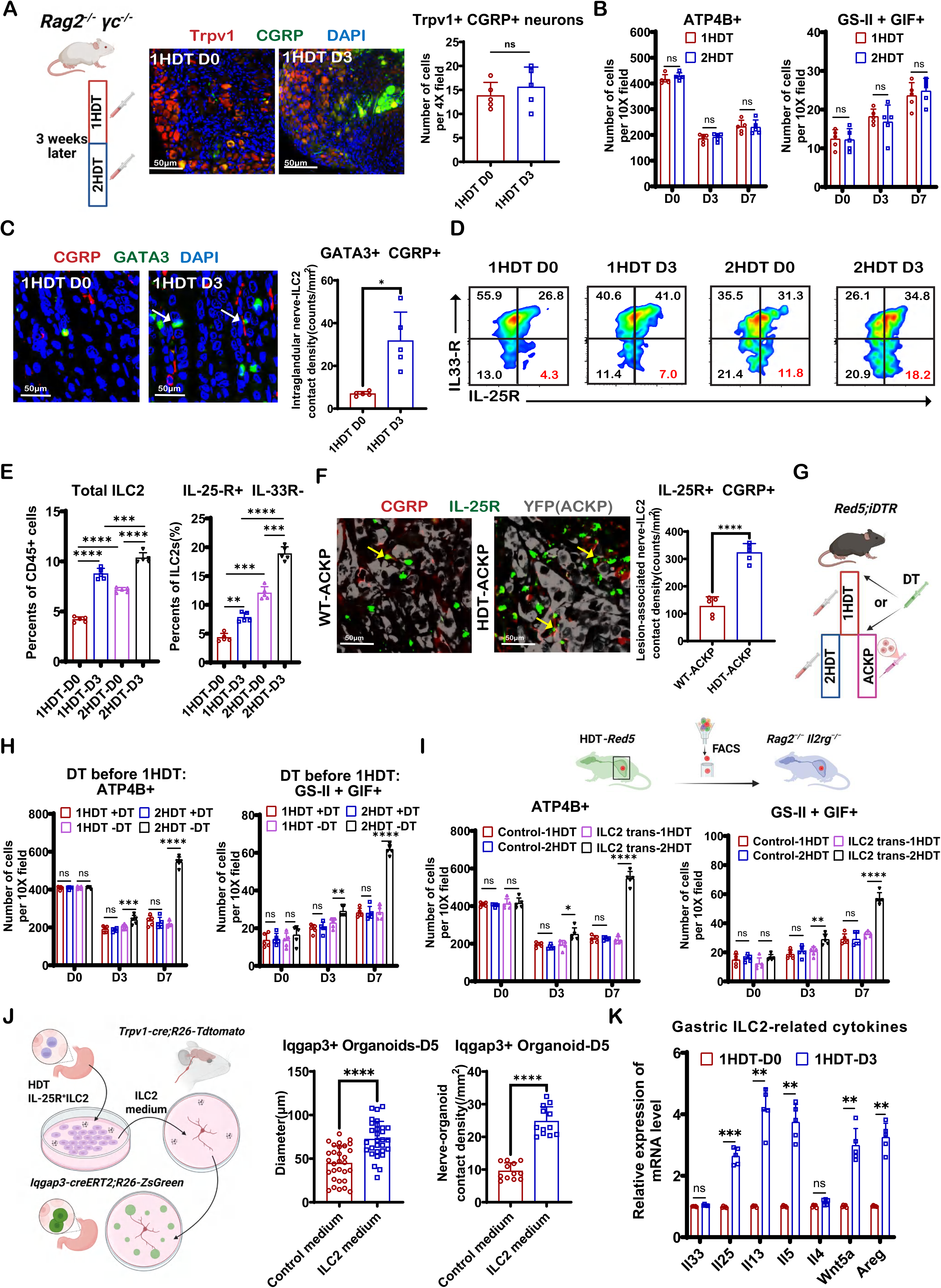
ILC2 signaling establishes an immune–sensory neuron circuit that reinforces gastric inflammatory memory (A, **B)** Representative immunofluorescence images and quantification of Trpv1^+^CGRP^+^ sensory neurons and gastric epithelial responses (ATP4B^+^ parietal cells and GSII^+^ GIF^+^ SPEM cells) in *Rag2^−/−^; Il2rg^−/−^* mice after primary injury (1HDT) or recall injury (2HDT) (n = 5 mice per group). Scale bars, 50 µm. **(C)** Representative immunofluorescence images showing GATA3^+^ gastric ILC2s and their spatial association with CGRP^+^ sensory fibers with quantification of ILC2–nerve contacts after HDT injury (n = 5 mice per group). Scale bars, 50 µm. **(D, E)** Representative flow cytometry plots and analysis of total gastric ILC2s and subsets (IL-25R^+^IL-33R^−^) (n = 5 mice per group). **(F)** Representative immunofluorescence Imaging and quantification of IL-25R^+^ cells in proximity to CGRP^+^ sensory fibers in ACKP gastric tumors (n = 5 mice per group). Scale bars, 50 µm. **(G)** Schematic of ILC2 ablation with different time point in *Red5; iDTR* mice. **(H)** Quantification of ATP4B⁺ cells, GSII⁺GIF⁺ cells with DT treatment before primary (1HDT) followed by recall (2HDT) injury in *Red5; iDTR* mice (n = 5 mice per group). **(I)** Schematic and quantification of ATP4B^+^ cells and GSII^+^GIF^+^ cells after adoptive transfer of ILC2s isolated from HDT-treated Red5 donor mice into *Rag2^−/−^; Il2rg^−/−^* recipients (n = 5 mice per group). **(J)** Schematic of ILC2–sensory neuron–organoid co-culture experiments with quantification of organoid diameter (n = 30 spheroids/group) and neuron–organoid contacts (n = 4 biologically independent mice per group, 3 fields analyzed per mouse). **(K)** Quantitative PCR analysis of gastric ILC2-associated cytokines at D0 and D3 after HDT (n = 5 mice per group). Statistical significance indicated by \**p* < 0.05, \*\**p* < 0.01, \*\*\**p* < 0.001, \*\*\*\**p* < 0.0001; ns, not significant. Data are mean ± s.e.m. *P* values were calculated using one-way ANOVA with Dunnett’s test (Figure 4H) or two-sided t-tests (Figure 4A**, 4B, 4C,4F, 4H,4I,4J and 4K)**. See also **Figure S7** and **S8**.

To characterize immune remodeling during gastric inflammatory recall, we performed flow cytometric analysis of gastric immune populations before and after HDT-induced injury **(Fig S7E)**. Total CD45⁺ immune cell infiltration was markedly increased during the acute inflammatory phase following HDT treatment **(Fig S7F)**. Among these populations, dendritic cells, macrophages, neutrophils, and innate lymphocytes exhibited the most prominent expansion, whereas T cells and monocytes showed comparatively modest changes **(Fig S7G)**. These findings indicate that recall injury is accompanied by robust innate immune activation within the gastric microenvironment. Among expanded immune populations, antibody-mediated depletion of innate lymphoid cells using anti-CD90.2 abolished the gastric memory phenotype, whereas depletion of other immune populations had no effect **(Fig S7H)**.

As type 2 innate lymphoid cells (ILC2s) constitute the major innate lymphoid population in the mouse stomach^22,39^, we focused on this subset. Spatial analysis revealed that following HDT injury, ILC2s preferentially localized to the gastric isthmus, with further enrichment during recall **(Fig S8A)**, and were found in close proximity to sensory nerve terminals **(Fig 4C)**, suggesting instructive interactions between ILC2s and sensory neurons. Flow cytometric analyses further showed that gastric ILC2s comprise two major subsets: natural ILC2s (nILC2s; IL-33R⁺ IL-25R⁻) and inflammatory ILC2s (iILC2s; IL-33R⁻ IL-25R⁺)^12,39,40^**(Fig 4D and S8B)**. Notably, the IL-25R⁺ iILC2 subset remained persistently abundant following injury and into the memory phase **(Fig 4E)**, indicating selective enrichment of this population, consistent with IL-25–biased and memory-like ILC2s reported previously^12^. This is in line with prior studies showing that HDT induces robust IL-25 production from gastric tuft cells, which preferentially activates and expands iILC2s^39^. In tumor models, IL-25R⁺ ILC2s were similarly enriched in mice with prior inflammation and displayed increased contacts with CGRP⁺ sensory terminals (Fig 4F).

To directly test the necessity of ILC2s, we selectively ablated ILC2s using *Red5; iDTR* mice^41^**(Fig 4G)**. Pre-depletion of ILC2s with diphtheria toxin before the initial HDT injury had minimal effects on baseline gastric architecture or the primary injury response, but completely abolished recall-associated regenerative acceleration during secondary injury **(Fig 4H)**. In contrast, depletion of ILC2s after the initial HDT episode failed to eliminate the memory phenotype **(Fig S8C)**, indicating that ILC2s are specifically required during the establishment phase, rather than the maintenance phase, of gastric inflammatory memory. Conversely, transplantation of activated ILC2s into *Rag2^⁻/⁻^γc^⁻/⁻^* mice restored the memory phenotype **(Fig 4I)**, establishing their sufficiency. Conditioned media from activated but not resting ILC2s primed nodose ganglia to enhance neuronal support of gastric organoid growth **(Fig 4J)**, indicating that ILC2s instruct neuronal memory via soluble factors.

Consistent with this role, qPCR analysis of sorted gastric ILC2s revealed increased expression of Il5, Il13, Wnt5a and Areg following injury **(Fig 4K)**. Although both Wnt5a and AREG have been implicated in axon growth^42,43^, only Wnt5a promoted NG axonal outgrowth **(Fig S8D)**, consistent with enrichment of its receptor FZD3—but not EGFR—in NG neurons based on analysis of a published single-cell RNA-seq dataset^44^ **(Fig S8E)**. In a three-way co-culture system, Wnt5a⁺/⁺ IL-25R⁺ ILC2s enhanced sensory neuron–mediated support of organoid growth and increased contacts between sensory nerve terminals and epithelial organoids, whereas this effect was attenuated with Wnt5a⁺/⁻ ILC2s **(Fig S8F)**, indicating that ILC2-derived Wnt5a facilitates neuro–epithelial interactions during memory establishment.

Finally, consistent with our previous work showing that isthmus localization of ILC2s depends on CXCL12/CXCR4 signaling^45^, conditional deletion of CXCR4 in ILC2s using *Red5; Cxcr4^flox/flox^*mice impaired their isthmus localization, reduced sensory innervation, and abolished inflammatory memory **(Fig S8G and S8H)**. Collectively, these results establish gastric ILC2s as the key innate immune population that instructs sensory neuron–mediated inflammatory memory.

### Epigenetic remodeling stabilizes inflammatory memory in sensory neurons

To elucidate how ILC2s “write” inflammatory memory into sensory neurons, we examined the impact of ILC2-derived cytokines on sensory neuronal function. Among cytokines released by HDT-activated ILC2s, IL-5 and IL-13 remained persistently elevated during the memory phase **(Fig S9A)**. Stimulation with IL-5 markedly enhanced CGRP release from sensory neurons, an effect significantly amplified in neurons isolated from mice with established inflammatory memory **(Fig 5A,5B and S9B, S9C)**, indicating that memory-encoded neurons exhibit heightened responsiveness to type 2 cytokine signaling.

**Figure 5.**
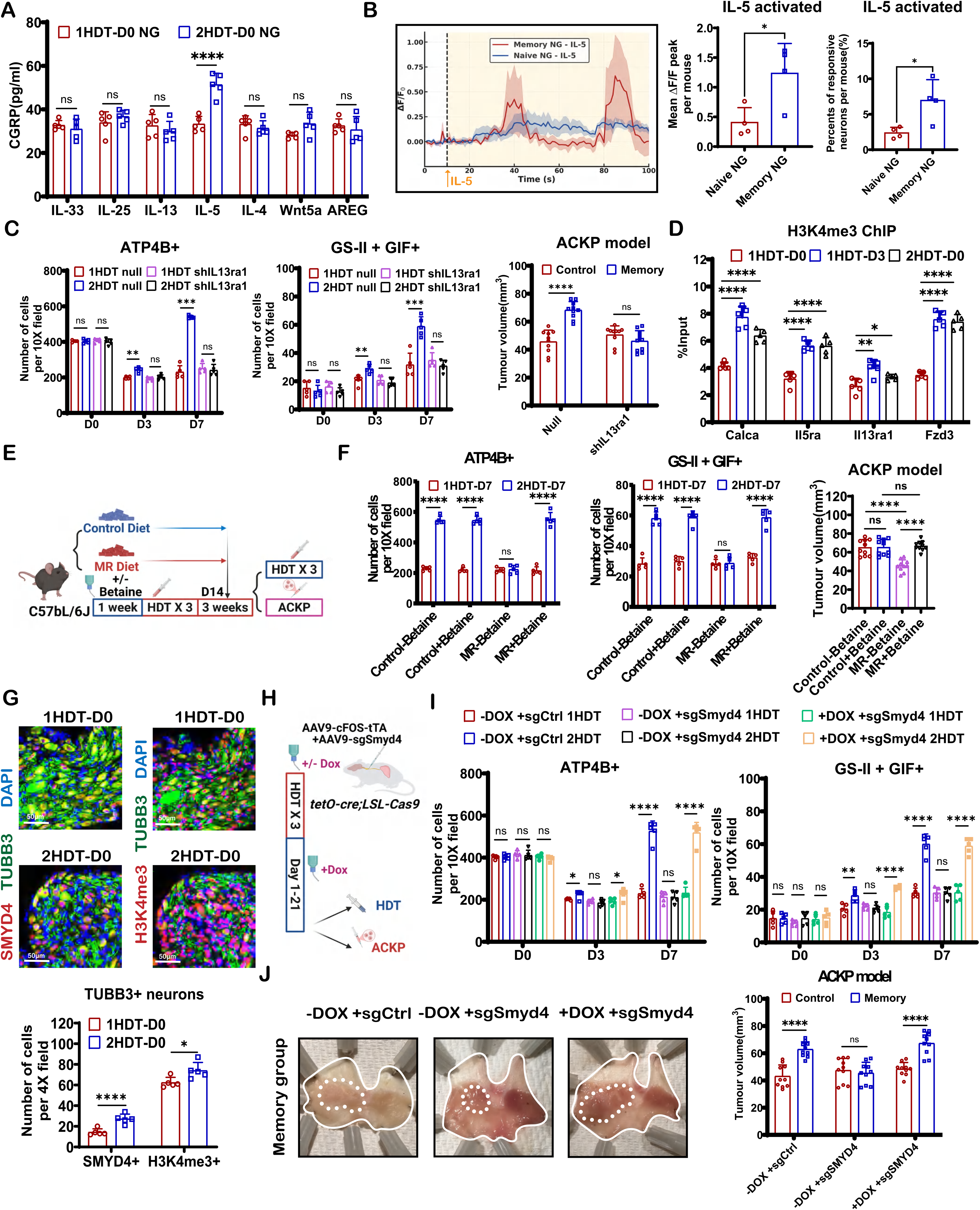
Epigenetic remodeling encodes long-term inflammatory memory in sensory neurons. **(A)** ELISA quantification of CGRP levels in nodose ganglia after 1HDT or 2HDT (n = 5 mice per group). **(B)** Nodose ganglia from *Trpv1-cre; R26-GCaMP6s* mice stimulated with IL-5 with fluorescence images and Ca²⁺ response traces with quantification from naive and inflammation-experienced mice (n = 5 mice per group). **(C)** Quantification of ATP4B⁺ cells and GSII⁺GIF⁺ cells (n = 5 mice per group) and tumor burden (n = 10 mice per group) in the orthotopic ACKP model with rAAV-shIl13ra1 or control into NG (n = 5 mice per group). **(D)** ChIP–qPCR analysis of H3K4me3 enrichment at promoters of indicated genes (n = 5 mice per group). **(E, F)** Methionine restriction or betaine supplementation followed by HDT injury and orthotopic ACKP tumor implantation with quantification of epithelial responses (n = 5 mice per group) and tumor volume (n = 10 mice per group). **(G)** TUBB3⁺ sensory neurons co-stained for SMYD4 or H3K4me3 with quantification of SMYD4⁺ and H3K4me3⁺ neurons before and after HDT injury (n = 5 mice per group). Scale bars, 50 µm. **(H, I)** Schematic of Fos-dependent targeting of SMYD4 using rAAV-cFos-tTA and rAAV-sgSmyd4 or control in *tetO-Cre; LSL-Cas9* mice with quantification of ATP4B⁺ and GSII⁺GIF⁺ cells after 1HDT or 2HDT (doxycycline on or off, n = 5 mice per group). **(J)** Representative gross images and quantification of tumor burden in the orthotopic ACKP model after Fos-targeted SMYD4 deletion under doxycycline-on or -off conditions (n = 10 mice per group). Statistical significance indicated by \**p* < 0.05, \*\**p* < 0.01, \*\*\**p* < 0.001, \*\*\*\**p* < 0.0001; ns, not significant. Data are mean ± s.e.m. *P* values were calculated using one-way ANOVA with Dunnett’s test (Figure 5D **and 5F)** or two-sided t-tests (Figure 5A**,5B,5C,5F,5G ,5I and 5J)**. See also **Figure S9** and **S10**.

Consistently, memory-encoded sensory neurons expressed higher levels of type 2 cytokine receptors, including IL-5RA, IL-13RA1, and FZD3, whereas IL-4R expression remained unchanged **(Fig S9D and S9E)**, revealing a long-lasting increase in neuronal cytokine sensitivity. Functional dissection of these pathways using either IL-5/IL-13 neutralizing antibodies during HDT treatment or targeted knockdown of IL5ra and IL13ra1 in NG neurons via AAV-shIL5ra and AAV-shIL13ra1 prior to HDT revealed distinct roles for these cytokine pathways. Disruption of IL-5 signaling partially attenuated CGRP release and recall responses but did not eliminate inflammatory memory, whereas inhibition of IL-13 receptor signaling completely abolished accelerated regeneration and tumor-promoting recall phenotypes **(Fig 5C and S9F–S9H)**. These findings define a cooperative cytokine circuit in which IL-5 amplifies neuronal output during recall, while IL-13 serves as the dominant signal responsible for writing and stabilizing inflammatory memory in sensory neurons.

Given the long-term persistence of inflammatory memory, we hypothesized that its maintenance relies on epigenetic remodeling rather than transient signaling. Accordingly, we observed enrichment of promoter-associated H3K4 trimethylation (H3K4me3) and enhancer-associated histone H3 lysine 27(H3K27) acetylation (H3K27ac) in the nodose ganglion during memory formation, whereas H3K4 monomethylation (H3K4me1) remained unchanged **(Fig S10A)**. Notably, these epigenetic alterations were largely abolished in NG neurons with IL13ra1 knockdown **(Fig S10B)**, indicating that IL-13 signaling serves as a major upstream input driving epigenetic remodeling during the establishment of sensory inflammatory memory. ChIP–qPCR analyses of Trpv1⁺ neurons revealed increased H3K4me3 and H3K27ac at regulatory regions of Calca, Il5ra, Il13ra1, and Fzd3 **(Fig 5D and S10C)**, linking chromatin remodeling to sustained CGRP production and cytokine receptor expression.

To assess the functional relevance of these histone modifications, we modulated neuronal methyl donor availability through dietary methionine restriction^46^ and betaine supplementation^47^. Although these interventions minimally affected baseline neuronal activity or acute injury responses, they effectively altered histone methylation in nodose ganglion neurons without affecting acetylation **(Fig S10D)**. Functionally, methionine restriction abolished, whereas betaine supplementation restored, accelerated regeneration and tumor-promoting recall responses **(Fig 5E and 5F)**. Notably, methionine restriction during the eradication phase of prior *H. pylori* infection similarly eliminated the inflammatory memory phenotype **(Fig S10E-S10G)**, indicating that H3K4me3 is a principal epigenetic mechanism governing inflammatory memory.

Mechanistically, analysis of a publicly available transcriptomic dataset of mouse nodose ganglia exposed to different cytokines^48^ identified SMYD4 as the only epigenetic regulator selectively induced by IL-13 stimulation **(Fig. S10H)**. Importantly, SMYD4 has been reported to function as an H3K4 trimethyltransferase^49^, nominating it as a compelling candidate linking IL-13 signaling to the persistent H3K4me3 remodeling observed during sensory inflammatory memory. Consistent with this possibility, SMYD4 expression remained persistently elevated in nodose ganglia during the memory phase, paralleling increased H3K4me3 enrichment **(Fig. 5G and S10I)**. Notably, persistent SMYD4 upregulation was abolished in NG neurons with IL13ra1 knockdown, indicating that IL-13 signaling is a dominant upstream driver of sustained SMYD4 activation during sensory inflammatory memory formation **(Fig S10J)**. To directly test the functional requirement for SMYD4, we employed an activity-dependent, neuron-specific CRISPR strategy to delete *Smyd4* in inflammation-activated sensory neurons **(Fig 5H)**. Although loss of SMYD4 did not affect baseline neuronal function, it prevented the establishment of H3K4me3 enrichment and completely abolished recall-associated accelerated regeneration, metaplasia, and tumor-promoting effects (Fig. 5I, J and S10K). Consequently, mice lacking neuronal SMYD4 exhibited significantly improved survival during gastric tumor progression (Fig. S10L). Together, these findings identify SMYD4 as a central epigenetic regulator of sensory neuronal inflammatory memory and suggest that disrupting memory maintenance may represent a therapeutic strategy to mitigate inflammation-driven cancer susceptibility.

## DISCUSSION

Our study identifies sensory neurons as a previously unrecognized component of long-term tissue memory in the stomach. Prior inflammatory injury establishes a durable neuronal state that can be reactivated upon recurrent tissue damage, thereby accelerating epithelial regeneration while simultaneously promoting metaplasia and tumor progression. This persistent program is encoded within Trpv1⁺ sensory neurons and maintained through IL-13–dependent SMYD4-mediated H3K4 trimethylation.

Inflammatory memory has thus far been studied primarily in epithelial and immune systems^6–12^. Epithelial stem and progenitor cells can retain epigenetic states associated with enhanced regeneration^8,9^, while innate immune populations may adopt trained inflammatory programs following prior stimulation^12^. Our findings extend this framework to the peripheral nervous system and suggest that tissue memory may emerge through coordinated adaptations across multiple cellular compartments rather than through a single autonomous cellular program. Importantly, these distinct systems may contribute different functions to long-term tissue adaptation. Epithelial cells may retain locally primed regenerative states, whereas immune populations provide inflammatory instruction and contextual signaling. However, many previously described inflammatory memory phenotypes remain highly dependent on intact tissue environments for their full manifestation. Although epithelial cells isolated from previously injured tissues can retain primed transcriptional or epigenetic states ex vivo^10,11^, enhanced regenerative responses are most prominently observed within multicellular tissue contexts^7^, suggesting that long-term tissue adaptation may require persistent support from additional systems capable of sustaining and recalling prior inflammatory experience.

Sensory neurons possess several unique properties that position them to fulfill such functions, including long lifespan, anatomical persistence, and circuit-level integration^14^. Unlike rapidly renewing epithelial populations, sensory neurons remain stably embedded within tissues and are capable of continuously integrating inflammatory information over extended periods^14^. Our findings therefore support a framework in which tissue memory emerges through coordinated interactions between epithelial, immune, and neural states, with sensory neurons providing a uniquely persistent substrate for long-term recall responses. A key feature of this neuronal state is its functional persistence and transferability. Memory-encoded neurons remain capable of driving epithelial responses long after tissue recovery, and nodose ganglion transplantation is sufficient to transfer enhanced regenerative phenotypes in vivo. These findings distinguish neuronal memory from transient inflammatory activation and support the idea that prior inflammatory experience is stably encoded within discrete sensory neuronal populations.

Our findings further identify ILC2s as critical immune instructors linking inflammatory context to neuronal memory establishment. While adaptive immunity is dispensable, ILC2 activation is both necessary and sufficient to establish the neuronal memory state. This instruction appears to operate through distinct but cooperative mechanisms. ILC2-derived Wnt5a promotes sensory axon extension and positioning within the gastric isthmus niche, whereas cytokines released by activated ILC2s directly reprogram neuronal states. Functional interrogation revealed a division of labor between these cytokine pathways. IL-5 primarily amplifies neuronal output during recall responses, whereas IL-13 acts as the dominant memory-writing signal responsible for establishing and stabilizing long-term neuronal adaptation. Through IL-13–dependent epigenetic remodeling, ILC2s induce stable neuronal reprogramming that persists long after resolution of inflammation. These findings extend recent observations of adaptive inflammatory states in innate lymphoid populations and position ILC2s as active participants in long-term tissue memory biology rather than transient inflammatory effectors.

At the molecular level, IL-13 signaling drives persistent SMYD4 upregulation and H3K4 trimethylation in sensory neurons. Notably, these epigenetic changes are restricted to neurons activated during the initial inflammatory episode, suggesting that inflammatory memory is encoded within a discrete neuronal ensemble rather than diffusely distributed throughout the sensory ganglion. This organization bears conceptual similarities to neuronal engrams described in learning and memory, raising the possibility that inflammatory experiences reshape neuronal identity through mechanisms analogous to those governing long-term information storage in the nervous system. During recall, memory-encoded sensory neurons translate stored inflammatory information into epithelial behavior through CGRP release and direct communication with gastric progenitor cells. Consistent with this model, pharmacological inhibition of CGRP–RAMP1 signaling suppresses regenerative and tumor-promoting recall responses and improves survival in memory-bearing tumor models, identifying CGRP as a key downstream effector through which neuronal memory influences tissue behavior.

Importantly, despite its long-term persistence, neuronal inflammatory memory remains at least partially reversible. Dietary methionine restriction effectively attenuated the memory phenotype without disrupting baseline neuronal function or acute injury responses, highlighting the plasticity of memory-encoded neuronal states. Likewise, loss of SMYD4 prevented the establishment and maintenance of the epigenetic landscape required for memory recall, resulting in suppression of regenerative, metaplastic, and tumor-promoting memory outputs. These observations suggest that neuronal inflammatory memory depends on active maintenance of a permissive chromatin state rather than representing a permanently fixed cellular program. Although methionine restriction constitutes a systemic metabolic intervention, our findings raise the possibility that selective targeting of SMYD4-dependent epigenetic mechanisms may provide a more precise therapeutic strategy for erasing maladaptive inflammatory memory while preserving normal tissue repair and neuronal function.

Finally, our findings provide a mechanistic explanation for the persistent elevation of gastric cancer risk following *H. pylori* eradication. By encoding prior inflammatory experience within sensory neurons, the nervous system establishes a long-lasting driver of epithelial hyper-responsiveness and tumor susceptibility. Collectively, our findings support a model in which inflammatory memory emerges through coordinated long-term adaptations across epithelial, immune, and neural compartments, with sensory neurons serving as a uniquely persistent substrate that links prior inflammatory experience to future tissue regeneration and cancer susceptibility.

### Limitations of the study

Several important questions remain unresolved. Although we demonstrate that peripheral inflammatory memory signals can propagate to central nervous system structures following gastric injury, it remains unclear whether the central nervous system itself subsequently undergoes long-term circuit remodeling capable of further modulating gastric responses upon recurrent injury. Thus, whether inflammatory memory ultimately becomes distributed across bidirectional gut–brain circuits remain to be determined. In addition, the organ specificity of sensory inflammatory memory remains unknown. While our findings establish a durable neuronal memory program in the stomach, it is unclear whether similar mechanisms operate across distinct tissues or whether inflammatory experience in one organ can influence future responses in distant organs through shared neural circuits. Addressing these questions will be important for understanding whether sensory inflammatory memory represents a localized tissue phenomenon or a broader systemic principle of neuroimmune adaptation.

## RESOURCE AVAILABILITY

### Lead contact

Information and requests for reagents may be directed to, and will be fulfilled by, the lead contact, Timothy C. Wang.

### Materials availability

Reagents are available on request from the lead contact with a completed materials transfer agreement. Information on reagents used in this study is available in the key resources table.

### Data and code availability

The scRNA-seq dataset for mouse JNCs was obtained from Kupari, J., et al., 2019^44^. The data are publicly available in Gene Expression Omnibus (GEO) at GSE124312 (https://www.ncbi.nlm.nih.gov/geo/query/acc.cgi?acc=GSE124312). The bulk RNA-seq dataset for mouse JNCs treated with different cytokines was obtained from Crosson, T., et al., 2024^48^. The data are publicly available in Gene Expression Omnibus (GEO) at GSE227968 (https://www.ncbi.nlm.nih.gov/geo/query/acc.cgi?acc=GSE227968).

## ACKNOWLEDGMENTS

This research was funded through grants from the NIH/NCI, including R01DK48077 and R35CA210088, as well as the Department of Defense grant W81XWH-21-1-0901 to T.C. Wang. This work was supported by Michael F. Price Memorial Grant Award from DeGregorio Family Foundation to T.C. Wang. This work was supported by NIH/NCI Cancer Center Support Grant P30CA013696. This work used Molecular Pathology/MPSR, Genomics and High Throughput Screening Shared Resource, Oncology Precision Therapeutics and Imaging Core (OPTIC) Shared Resource which is supported by funds from the Columbia University Medical Center Cancer Center Support Grant (CCSG) and NIH grant P30CA013696 (National Cancer Institute), and the resources of the Cancer Center Flow Core Facility funded in part through Grant P30CA013696. This study was funded in part through the NIH/NIDDK Columbia University Digestive and Liver Disease Research Center grant 5P30DK132710 and used the Bioimaging Core, Organoid & Cell Culture Core, and Clinical Biospecimen and Research Core. Imaging was performed with support from the Zuckerman Institute’s Cellular Imaging platform, funded through NIH Grant 1S10OD023587-01.

## AUTHOR CONTRIBUTIONS

Y.Z. and T.C.W. conceived and designed the study. Y.Z. performed most experiments and analyses. Y.Z. and P.Z. collaborated on calcium imaging and confocal microscopy experiments. Y.Z., F.W., J.Q., S.L and Q.W. performed flow cytometry experiments and contributed to data analysis strategies. R.T., M.H, J.L, J.A. and L.Z. assisted with animal experiments. X.Z. provided guidance on chemogenetic and optogenetic activation experiments. H.K. assisted with histopathological grading of mouse tissue sections. Y.O. provided imaging support for whole-mount staining. B.Z. assisted with the mouse ulcer model. H.Z. assisted with RNAscope experiments. T.C.W. supervised the project. Y.Z. and T.C.W. wrote the manuscript. All authors contributed substantially to the discussion of content for the article, reviewed and/or edited the manuscript prior to submission.

We acknowledge support from Columbia University shared resources and thank Dajiang “Kevin” Sun and Tingting Sun (Molecular Pathology/MPSR, Columbia University) for their expertise and assistance during this project. We thank Dr. Yanping Sun for her contributions in MRI data acquisition and analysis. We also thank Dr. Theresa Swayne and Dr. Yanyan Chen for their contributions in fluorescence imaging of live animal cells.

---

**Figure S1. Prior gastric inflammation establishes long-lasting recall responses across injury and regeneration models, related to Figure 1**

**(A)** Representative hematoxylin and eosin (H&E)-stained gastric sections at the indicated time points following primary high-dose tamoxifen injury (1HDT) or recall injury (2HDT). Scale bars, 100 µm.

**(B)** Representative immunofluorescence images showing CD45⁺ immune cell infiltration in gastric glands after 1HDT or 2HDT. Scale bars, 50 µm.

**(C)** Quantification of CD45⁺ immune cells in the gastric isthmus after 1HDT or 2HDT (n = 5 mice per group).

**(D)** Experimental schematic illustrating the HDT-induced injury, recovery and different recall timeline.

**(E)** Quantification of ATP4B⁺ cells, GIF⁺ cells and GSII⁺GIF⁺ cells after 1HDT or 2HDT of different recall timeline (n = 5 mice per group).

**(F)** Quantitative PCR analysis confirming successful *H.pylori(HP)* eradication following antibiotic treatment (n = 5 mice per group).

**(G)** Quantification of glandular epithelial lineage changes associated with *H. pylori* infection and after antibiotic eradication (n = 5 mice per group).

**(H)** Representative immunofluorescence images on day 7 after HDT challenge in control versus *H. pylori*-eradicated mice. Scale bars, 50 µm.

**(I)** Experimental schematics of the acetic acid-induced gastric ulcer healing model in inflammation-naive mice versus mice with prior HDT injury or *H. pylori* infection/eradication (ER).

**(J)** Representative gross images and quantification of ulcer area on day 7 after acetic acid-induced injury in the indicated groups (n = 5 mice per group).

Statistical significance indicated by \**p* < 0.05, \*\**p* < 0.01, \*\*\**p* < 0.001, \*\*\*\**p* < 0.0001; ns, not significant. Data are mean ± s.e.m. *P* values were calculated using one-way ANOVA with Dunnett’s test **(Figure S1C, S1E and S1F)** or two-sided t-tests **(Figure S1J)**.

**Figure S2. Prior gastric inflammation accelerates tumor progression in different cancer models, related to Figure 1**

**(A)** Experimental schematic for evaluating tumor progression in an inflammation-experienced diffuse-type gastric cancer model (*Mist1-CreERT2; CDH1^flox/flox^; RhoA^Y42C^*).

**(B)** Representative gross images of stomachs from control and *H. pylori*–eradicated mice in the diffuse-type gastric cancer model, with quantification of tumor volume (n = 10 mice per group).

**(C)** Representative H&E-stained gastric sections from control and *H. pylori*–eradicated mice with invasive tumor regions outlined. Right, quantification of invasion depth (shallow versus deep). Scale bars, 250 µm.

**(D)** Kaplan–Meier survival curves of control and *H. pylori*–eradicated mice in the diffuse-type gastric cancer model (n = 10 mice per group).

**(E)** Experimental schematic. *Iqgap3-CreERT2; R26-ZsGreen* mice were infected with *H. pylori*, followed by eradication. Three months later, mice received tamoxifen (TAM) to label Iqgap3⁺ gastric progenitor cells, which were isolated by FACS and subjected to organoid culture.

**(F)** Representative images and quantification of organoids derived from FACS-isolated Iqgap3⁺ (ZsGreen⁺) gastric progenitor cells from control and *H. pylori*-eradicated mice at days 5 and 10 of culture (n = 30 spheroids/group). Scale bars, 100 µm.

Statistical significance indicated by \**p* < 0.05, \*\**p* < 0.01, \*\*\**p* < 0.001, \*\*\*\**p* < 0.0001; ns, not significant. Data are presented as mean ± SEM. Two-sided unpaired Student’s t-test. Scale bars, 100 μm. Data are mean ± s.e.m. *P* values were calculated using two-sided t-tests **(Figure S2B and S2F)**, Fisher’s exact test **(Figure S2C)** or log-rank test **(Figure S2D)**.

**Figure S3. Sensory CGRP⁺ innervation is selectively enhanced in inflammation-associated regeneration and tumor contexts, related to Figure 2**

**(A)** Representative immunofluorescence images and quantification of CGRP⁺ sensory nerve density in gastric tissues from control and *H. pylori*–eradicated mice after HDT (n = 5 mice per group). Scale bars, 50 µm.

**(B-C)** Representative immunofluorescence images and quantification of TH⁺, ChAT⁺ and Substance P+ nerve density in gastric tissues after primary (1HDT) or recall (2HDT) injury (n = 5 mice per group). Scale bars, 100 µm.

**(D)** Representative immunofluorescence images and quantification of CGRP⁺ sensory nerve density in gastric tissues from control and HDT–treated mice after acetic acid–induced ulcer (n = 5 mice per group). Scale bars, 50 µm.

**(E-G)** Representative immunofluorescence images and quantification of CGRP⁺ sensory nerve density in the lesions from orthotopic ACKP model, intestinal-type gastric cancer model (*Iqgap3-CreERT2; LSL-Kras^G12D^* mice) and diffuse-type gastric cancer (*Mist1-CreERT2; CDH1^flox/flox^; RhoA^Y42C^* mice) (n = 5 mice per group). Scale bars, 100 µm (S3F) and 50 µm (S3E and S3G).

**(H)** Quantification of c-Fos⁺CGRP⁺ and Trpv1⁺CGRP⁺ neurons in nodose ganglia after 1HDT or 2HDT (n = 5 mice per group).

**(I, J)** Representative immunofluorescence images and quantification of nodose ganglia stained for Trpv1 and CGRP in mice bearing orthotopic ACKP tumors after HDT treatment or control (n = 5 mice per group). Scale bars, 100 µm.

**(K)** Representative gross images showing the stomach before and after UVT.

**(L)** Representative immunofluorescence images and quantification of CGRP⁺ sensory nerve density in the anterior versus posterior gastric wall after HDT following UVT (n = 5 mice per group). Scale bars, 50 µm.

**(M)** Representative immunofluorescence images and quantification of ATP4B⁺ cells and GSII⁺GIF⁺ cells after recall injury(2HDT) from UVT mice or control (n = 5 mice per group). Scale bars, 50 µm.

**(N)** Representative gross images and quantification of tumor burden in orthotopic ACKP tumors after HDT in control and UVT-treated mice (n = 10 mice per group).

Statistical significance indicated by \**p* < 0.05, \*\**p* < 0.01, \*\*\**p* < 0.001, \*\*\*\**p* < 0.0001; ns, not significant. Data are mean ± s.e.m. Statistical analyses were performed using two-sided t-tests.

**Figure S4. Gastric inflammatory memory propagates to central vagal circuits and is encoded in sensory ganglia, related to Figure 2**

**(A)** Brainstem MRI images showing diffusion-weighted sequences with regions of interest corresponding to the nucleus tractus solitarius (NTS).

**(B)** Quantification of diffusion coefficient (ADC), T1 and T2 signals in the NTS (n = 3 mice per group).

**(C)** c-Fos⁺ neurons immunofluorescence staining and quantification in the NTS after HDT injury (n = 5 mice per group). Scale bars, 50 µm.

**(D)** Representative immunofluorescence images and quantification of ATP4B⁺ and GSII⁺GIF⁺ cells after HDT injury in *Trpv1-Cre; R26-hM3Dq* mice with Clozapine-N-oxide (CNO) treatment (n = 5 mice per group). Scale bars, 50 µm.

**(E)** Experimental schematic and quantification of ATP4B⁺ and GSII⁺GIF⁺ cells of repeated HDT injury in *Trpv1-Cre; R26-DTA* mice (n = 5 mice per group).

**(F)** Representative gross images of ACKP tumor from *tetO-Cre; R26-tdTomato; R26-DTA* mice under doxycycline-on conditions, followed by HDT injury or control.

**(G-H)** Gross images and H&E sections showing nodose ganglia transplanted into the gastric subserosa. H&E sections, Scale bars, 100 µm.

**(I)** Quantification of transplanted nodose ganglion (NG) graft survival and axonal outgrowth in recipient mice.

**(J)** Representative immunofluorescence images of gastric tissues following memory NG transplantation stained for tdTomato, p-ERK, and DAPI. White lines indicate the anatomical boundaries of the submucosa and subserosa. Scale bars, 100 μm.

**(K)** ATP4B and GIF/GSII immunofluorescence staining in gastric glands surrounding transplanted ganglia from memory group or control (n = 5 mice per group). Scale bars, 50 µm.

**(L)** Gross images of tumor burden in mice transplanted with control or memory-experienced ganglia (n = 10 mice per group).

Statistical significance indicated by \**p* < 0.05, \*\**p* < 0.01, \*\*\**p* < 0.001, \*\*\*\**p* < 0.0001; ns, not significant. Data are mean ± s.e.m. Statistical analyses were performed using two-sided t-tests**(Figure S4B, S4C, S4D, S4E and S4I)** or Fisher’s exact test **(Figure S4I)**.

**Figure S5. CGRP/Ramp1 signaling is sufficient to promote gastric epithelial activation and regeneration, related to Figure 3**

**(A)** Representative immunofluorescence ATP4B and GIF/GSII staining images showing the effects of exogenous CGRP administration in *Rag2^−/−^; Il2rg^−/−^* mice subjected to repeated HDT injury. Scale bars, 50 µm.

**(B)** Representative immunofluorescence images showing Ki67 staining in gastric glands after HDT in saline- or CGRP-treated mice. Scale bars, 50 µm.

**(C)** Experimental schematic and quantification of ATP4B⁺ cells, GSII⁺GIF⁺ cells and Ki67⁺ cells (n = 5 mice per group).

**(D)** Representative RNAscope images and quantification of Iqgap3 and Ramp1 mRNA expression in gastric isthmus region at different time points (n = 5 mice per group).

**(E)** Quantification of ATP4B⁺ cells and GS-II⁺GIF⁺ cells in control and Ramp1 conditional knockout mice following HDT-induced gastric injury with saline or CGRP administration (n = 5 mice per group).

**(F)** Representative images showing ATP4B and GIF/GSII staining in gastric glands from *Iqgap3-CreERT2; Ramp1^flox/flox^* mice (n = 5 mice per group). Scale bars, 50 µm.

**(G)** Bright-field images of gastric organoids cultured with or without nodose ganglia. Scale bars, 50 µm.

**(H)** Experimental schematic illustrating nodose ganglion–organoid co-culture.

**(I)** Representative fluorescence images showing Trpv1-lineage sensory neurons and Iqgap3⁺ gastric organoids with quantification of organoid diameter (n = 30 organoids per condition). Scale bars, 100 µm.

Statistical significance indicated by \**p* < 0.05, \*\**p* < 0.01, \*\*\**p* < 0.001, \*\*\*\**p* < 0.0001; ns, not significant. Data are mean ± s.e.m. *P* values were calculated using one-way ANOVA with Dunnett’s test **(Figure S5D and S5I)** or two-sided t-tests **(Figure S5C and S5E)**.

**Figure S6. Synaptic CGRP release from sensory neurons is required for epithelial activation and tumor promotion, related to Figure 3**

**(A)** Whole-mount fluorescence image showing CGRP⁺ sensory nerve fibers in proximity to Iqgap3⁺ cells in the gastric isthmus. Below, three-dimensional reconstruction of the indicated glandular region. Scale bars, 100 µm.

**(B)** Representative immunofluorescence images and quantification showing synaptophysin-labelled nerve fibers associated with Iqgap3⁺ cells. Scale bars, 50 µm (n = 5 mice per group).

**(C)** Experimental schematic of AAV9-hSyn-FLEX-TeLC injection into the nodose ganglia of Trpv1-Cre mice. Representative immunofluorescence staining images with quantification of nodose ganglia collected on day 3 after the first (1HDT-D3) or second (2HDT-D3) gastric injury (n = 5 mice per group). Scale bars, 100 µm.

**(D)** Representative immunofluorescence images with quantification of gastric tissues from control and AAV-TeLC-treated mice after the gastric injury, stained for VAMP2, CGRP, and DAPI (n = 5 mice per group). Scale bars, 50 µm.

**(E)** Quantification of ATP4B⁺ cells and GSII⁺GIF⁺ cells following AAV-TeLC treatment or control (n = 5 mice per group).

**(F)** Experimental schematic illustrating pharmacological inhibition of the CGRP pathway using rimegepant.

**(G-H)** Representative images and quantification of ATP4B⁺ cells and GSII⁺GIF⁺ cells after repeated HDT injury with rimegepant treatment (n = 5 mice per group). Scale bars, 50 µm.

**(I)** Representative gross images and quantification of ACKP tumor volume with saline or rimegepant treatment after HDT injury (n = 10 mice per group).

**(J)** Kaplan–Meier survival curves of mice bearing orthotopic ACKP tumors treated with saline or rimegepant after HDT injury (n = 10 mice per group).

Statistical significance indicated by \**p* < 0.05, \*\**p* < 0.01, \*\*\**p* < 0.001, \*\*\*\**p* < 0.0001; ns, not significant. Data are mean ± s.e.m. Statistical analyses were performed using two-sided t-tests **(Figure S6B, S6C, S6D, S6E, S6I and S6H)** or log-rank test**(Figure S6J)**.

**Figure S7. Recall sensory neuron activation is not driven by broad immune cell composition changes, related to Figure 4**

**(A)** Experimental schematic and representative immunofluorescence images showing Trpv1/CGRP sensory neuron staining in *Rag1^−/−^* mice after HDT injury with quantification of Trpv1⁺CGRP⁺ neurons (n = 5 mice per group). Scale bars, 50 µm.

**(B)** Representative immunofluorescence images and quantification of CGRP⁺ sensory nerve density in gastric tissues after primary (1HDT) or recall (2HDT) injury in *Rag1^−/−^* mice (n = 5 mice per group). Scale bars, 50 µm.

**(C)** Representative images showing ATP4B⁺ cells and GSII⁺GIF⁺ cells in gastric glands after 1HDT or 2HDT with quantification of ATP4B⁺ and GSII⁺GIF⁺ cells in *Rag1^−/−^*mice (n = 5 mice per group). Scale bars, 50 µm.

**(D)** Representative images and quantification of CGRP⁺ sensory nerve density in *Rag2^−/−^; Il2rg^−/−^* mice after 1HDT or 2HDT (n = 5 mice per group). Scale bars, 50 µm.

**(E)** Representative flow cytometry plots showing major immune cell populations within CD45⁺ gastric immune cells.

**(F)** Quantification of total CD45⁺ immune cells after HDT injury (n = 5 mice per group).

**(G)** Quantification of immune cell subsets among CD45⁺ cells including T cells, dendritic cells, macrophages, neutrophils, monocytes and innate lymphocytes (n = 5 mice per group).

**(H)** Effects of immune cell subsets depletion on epithelial responses following primary or recall injury with quantification of ATP4B⁺ cells and GSII⁺GIF⁺ cells (n = 5 mice per group). Statistical significance indicated by \**p* < 0.05, \*\**p* < 0.01, \*\*\**p* < 0.001, \*\*\*\**p* < 0.0001; ns, not significant. Data are mean ± s.e.m. Statistical analyses were performed using two-sided t-tests.

**Figure S8. ILC2–sensory neuron interactions are mediated by Wnt5a signaling and defined by a distinct gastric ILC2 subset, related to Figure 4**

**(A)** Representative immunofluorescence images of gastric tissues collected before and after gastric injury (D3), stained for GATA3, CD3, and DAPI. Right, quantification of gastric isthmus-associated GATA3⁺CD3⁻ ILC2s following the first (1HDT) and second (2HDT) injury (n = 5 mice per group).

**(B)** Flow cytometric gating strategy used to identify gastric ILC2s.

**(C)** Quantification of ATP4B⁺ cells, GSII⁺GIF⁺ cells after primary or recall injury in mice with or without ILC2 ablation after first injury (n = 5 mice per group).

**(D)** Representative images of nodose ganglia stained for TUBB3 following treatment with NGF, Wnt5a or AREG with quantification of axonal outgrowth index (n = 5 ganglia per condition). Scale bars, 100 µm.

**(E)** Dot plots showing expression of candidate ligand–receptor pairs involved in ILC2–neuron communication, derived from the nodose ganglia single-cell RNA-seq dataset (GSE124312). Box plots show median (center line), interquartile range (box limits), and whiskers indicate 1.5× the interquartile range. Individual data points are shown.

**(F)** Experimental schematic of ILC2–organoid co-culture assays with representative images showing Trpv1-lineage sensory neurons and ZsGreen⁺ gastric epithelial cells. Quantification of organoid diameter (n = 30 organoids per condition) and nerve–organoid contact density (n = 4 biologically independent mice per group, 3 fields analyzed per mouse) is shown.

**(G)** Quantification of IL-25R⁺ gastric ILC2s and CGRP⁺ sensory nerve fibers in *Red5; Cxcr4^+/+^*or *Red5; Cxcr4^flox/flox^* mice followed by HDT-induced injury (n = 5 mice per group).

**(H)** Quantification of ATP4B⁺ cells and GSII⁺GIF⁺ cells following 1HDT or 2HDT in *Red5; Cxcr4^flox/flox^* or *Red5; Cxcr4^+/+^* mice (n = 5 mice per group).

Statistical significance indicated by \**p* < 0.05, \*\**p* < 0.01, \*\*\**p* < 0.001, \*\*\*\**p* < 0.0001; ns, not significant. Data are mean ± s.e.m. Statistical analyses were performed using one-way ANOVA **(Figure S8D and S8F)** or two-sided t-tests **(Figure S8A, S8C, S8G and S8H)**.

**Figure S9. IL-13 signaling enhances sensory neuron responsiveness during inflammatory recall, related to Figure 5**

**(A)** Representative flow cytometry plots and quantification showing IL-5⁺ and IL-13⁺ gastric ILC2s among IL-25R⁺ cells after primary or recall injury (n = 5 mice per group).

**(B, C)** Representative calcium imaging and traces showing nodose ganglion neuron responses to IL-5, IL-13, Wnt5a, AREG or capsaicin stimulation (n = 4 mice per group).

**(D)** Quantitative PCR analysis of cytokine and receptor gene expression in nodose ganglia after HDT injury (n = 5 mice per group).

**(E)** Representative images showing CGRP⁺ sensory neurons co-stained for IL5Rα, IL4Rα, IL13Rα1 or FZD3 with quantification (n = 5 mice per group). Scale bars, 50 µm.

**(F)** Quantification of ATP4B⁺ and GSII⁺GIF⁺ cells in mice treated with IL-5 neutralizing antibody or saline (n = 5 mice per group).

**(G)** Quantification of ATP4B⁺ parietal cells and GSII⁺GIF⁺ SPEM cells following nodose ganglion injection of rAAV-shIL5ra or control (n = 5 mice per group).

**(H)** Quantification of ATP4B⁺ parietal cells and GSII⁺GIF⁺ cells in mice treated with IL-13 neutralizing antibody or saline (n = 5 mice per group).

Statistical significance indicated by \**p* < 0.05, \*\**p* < 0.01, \*\*\**p* < 0.001, \*\*\*\**p* < 0.0001; ns, not significant. Data are mean ± s.e.m. Statistical analyses were performed using one-way ANOVA **(Figure S9A)** or two-sided t-tests **(Figure S9D, S9E, S9F, S9G and S9H)**.

**Figure S10. Sensory neuron inflammatory memory is associated with SMYD4-dependent H3K4me3 remodeling, related to Figure 5**

**(A)** Representative immunofluorescence images showing H3K4me3, H3K27ac and H3K4me1 staining in nodose ganglia before and after HDT injury with quantification (n = 5 mice per group). Scale bars, 50 µm.

**(B)** Quantification showing H3K4me3, H3K27ac and H3K4me1 staining in nodose ganglia with AAV-shIL13ra1 treatment or control before and after HDT injury (n = 5 mice per group).

**(C)** ChIP–qPCR analysis of H3K27ac enrichment at promoters of indicated genes in nodose ganglia before and after HDT injury (n = 5 mice per group).

**(D)** Quantification of circulating plasma S-adenosylmethionine (SAM) and H3K4me3 or H3K27ac modification–positive sensory neurons in mice subjected to control diet, methionine-restricted (MR) diet or betaine supplementation (n = 5 mice per group).

**(E)** Experimental schematic of *H.pylori* infection, eradication and methionine-restricted (MR) dietary intervention followed by HDT recall or orthotopic ACKP tumor.

**(F)** Quantification of H3K4me3⁺ neurons in nodose ganglia from control and previously H. pylori-infected mice maintained on control or methionine-restricted (MR) diets.

**(G)** Quantification of ATP4B⁺ cells, GS-II⁺GIF⁺ cells (n = 5 mice per group), and ACKP tumor volume (n = 10 mice per group) in control and previously *H. pylori*-infected mice maintained on control or methionine-restricted (MR) diets.

**(H)** Heat maps show transcriptional responses of nodose ganglia to IL-13 stimulation from dataset GSE227968.

**(I)** Representative immunofluorescence images showing SMYD4 and TUBB3 co-staining in nodose ganglia with quantification of SMYD4⁺ TUBB3⁺ neurons (n = 5 mice per group). Scale bars, 100 µm.

**(J)** Quantification showing SMYD4 and TUBB3 co-staining in nodose ganglia with AAV-shIL13ra1 treatment or control (n = 5 mice per group). Scale bars, 100 µm.

**(K)** Quantification of SMYD4⁺ and H3K4me3⁺ neurons following Fos-dependent SMYD4 deletion under doxycycline-on or -off conditions (n = 5 mice per group).

**(L)** Kaplan–Meier survival curves of mice bearing orthotopic ACKP tumors with Fos-dependent SMYD4 deletion under doxycycline-on or -off conditions (n = 10 mice per group). Statistical significance indicated by \**p* < 0.05, \*\**p* < 0.01, \*\*\**p* < 0.001, \*\*\*\**p* < 0.0001; ns, not significant. Data are mean ± s.e.m. Statistical analyses were performed using one-way ANOVA **(Figure S10C, S10D, S10F, S10I and S10K)**, two-sided t-tests **(Figure S10A, S10B, S10D, S10G and S10J)** or log-rank test**(Figure S6L)**.

## STAR★METHODS

### KEY RESOURCES TABLE

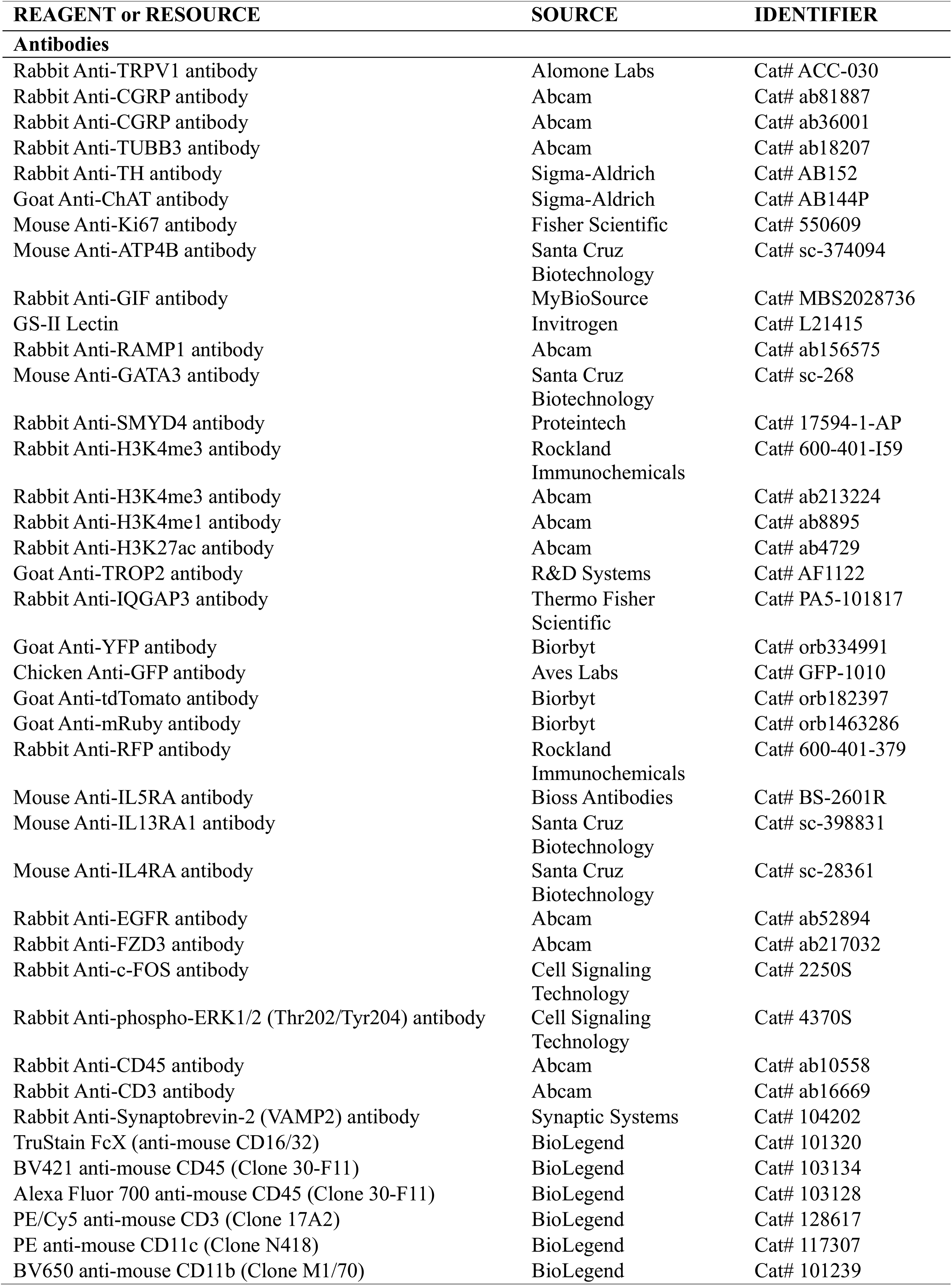
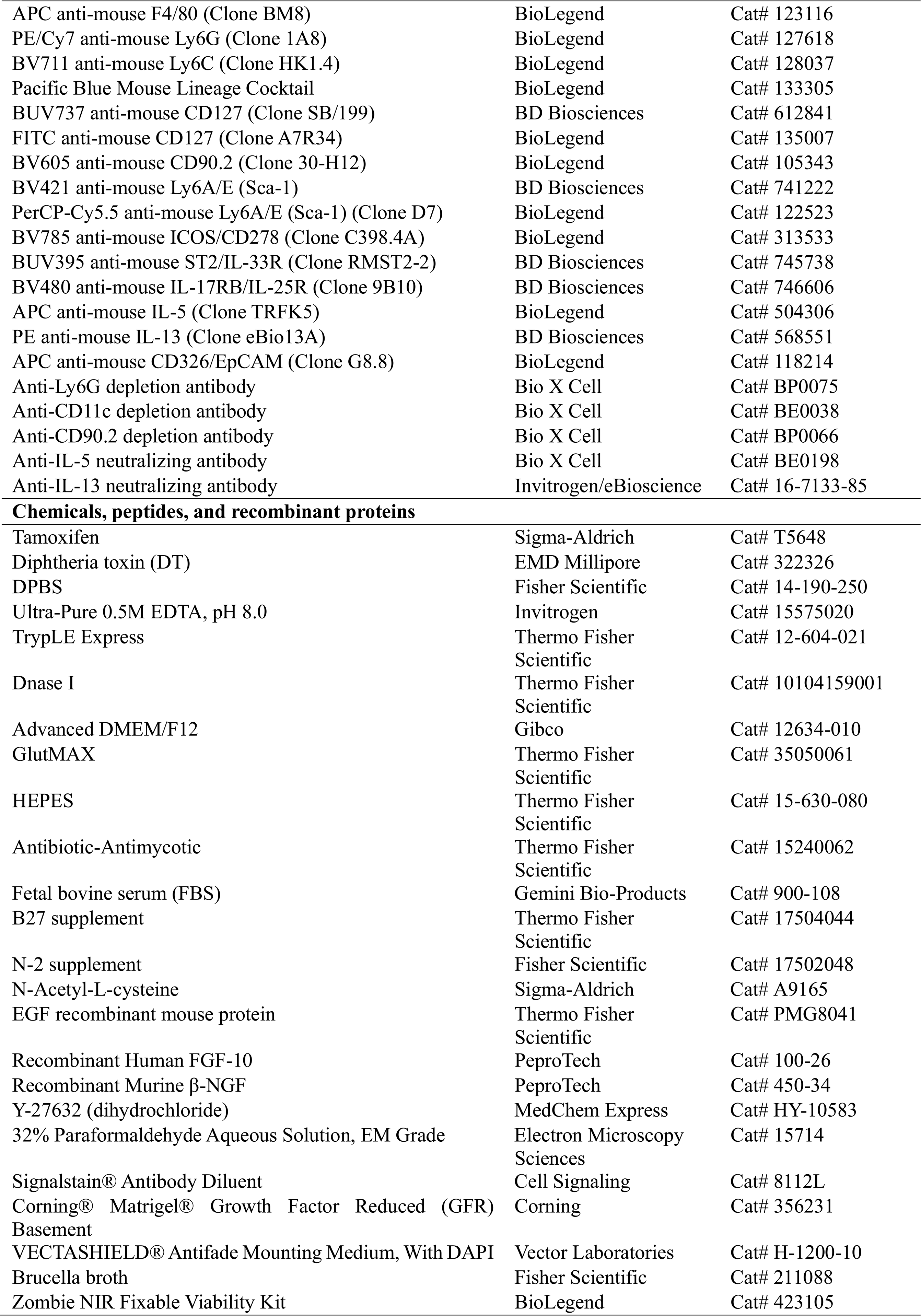
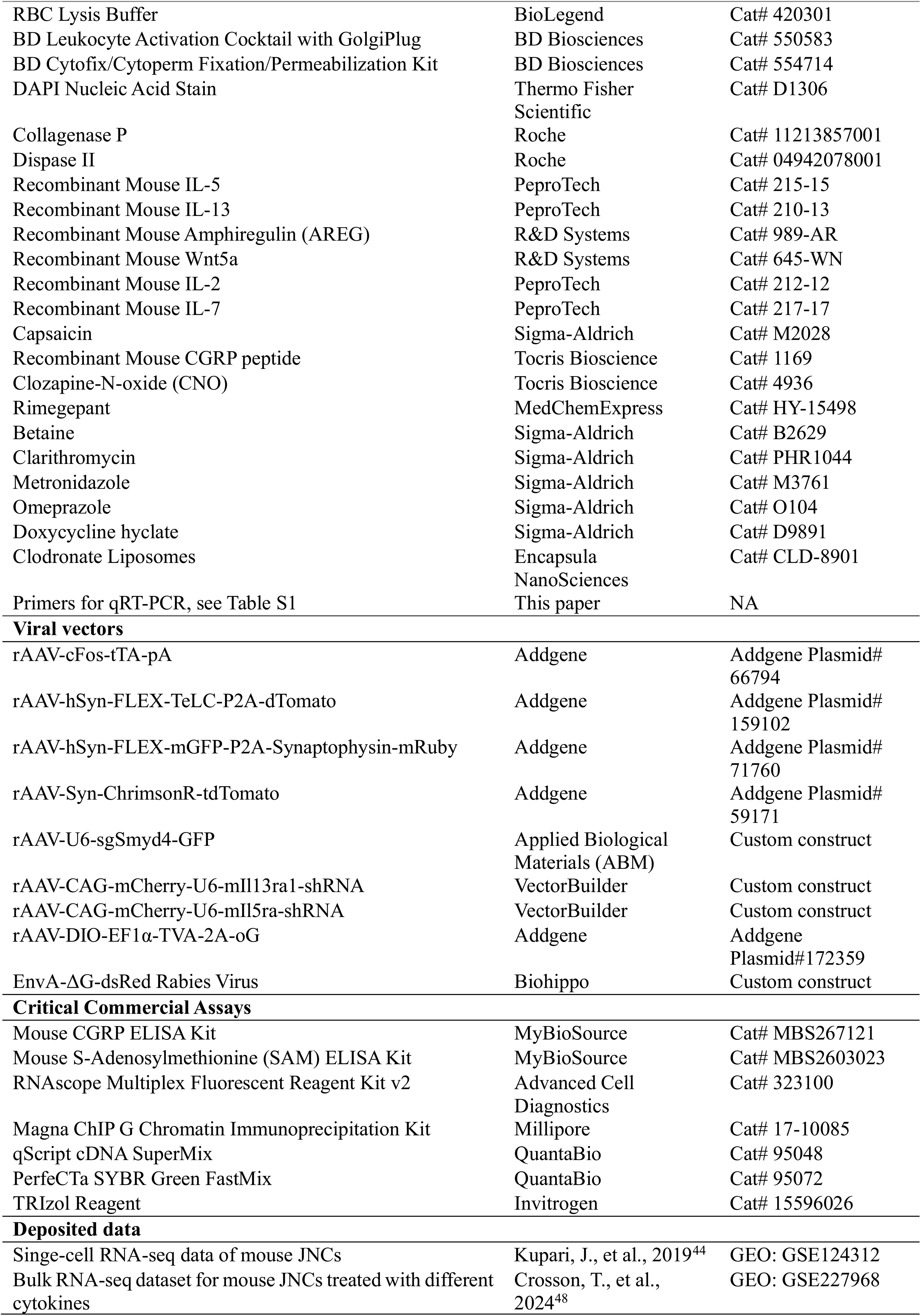
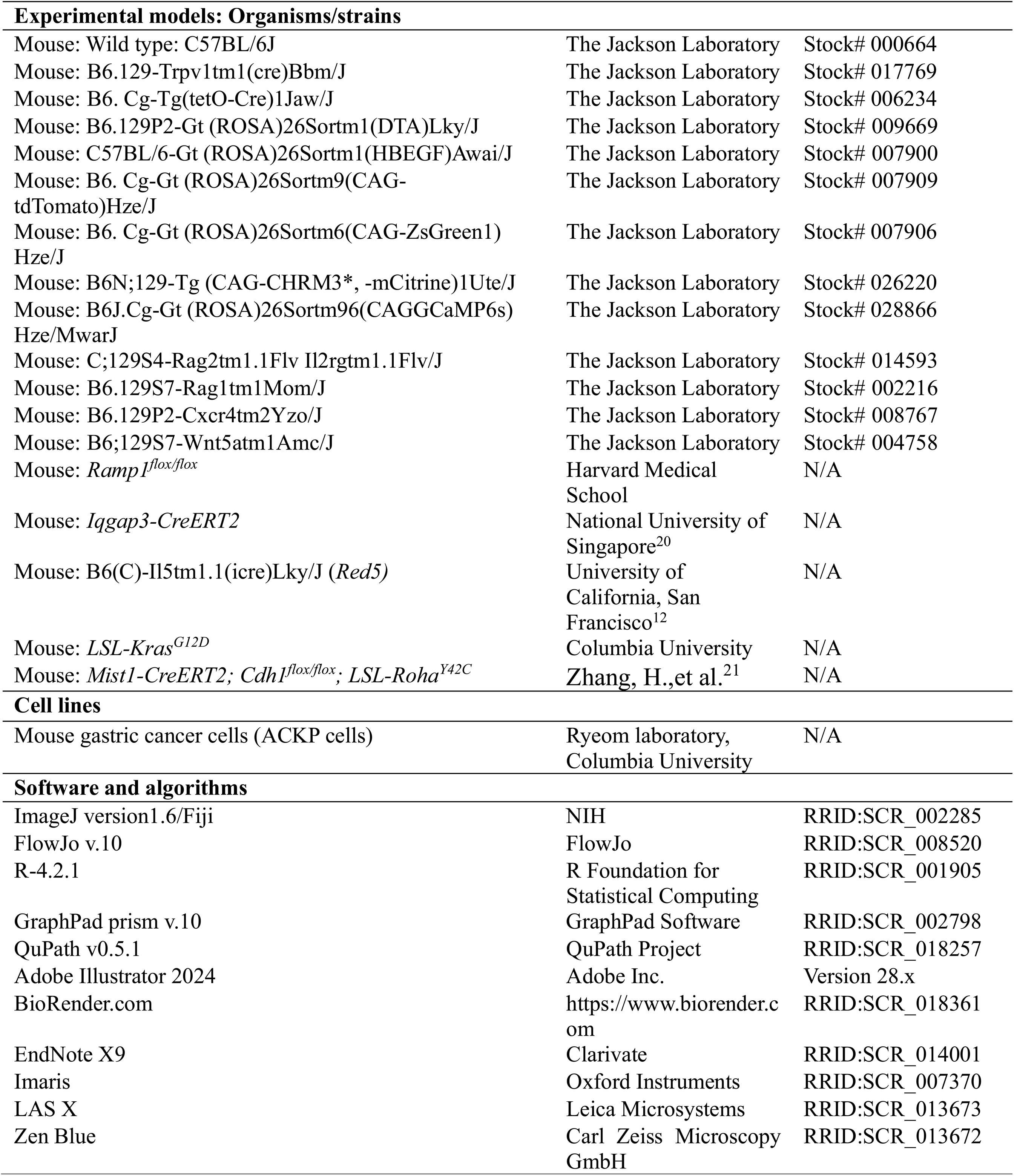

### EXPERIMENTAL MODEL AND SUBJECT DETAILS

#### Animals

All animal experiments were conducted in accordance with National Institutes of Health guidelines and were approved by the Institutional Animal Care and Use Committee (IACUC) of Columbia University.

Mice were maintained under specific pathogen-free conditions on a 12-h light–dark cycle (lights on, 07:00–19:00) at 22 °C and 40–50% humidity, with ad libitum access to food and water. Unless otherwise specified, experiments were performed in 8–12-week-old mice. Both male and female mice were used and age- and sex-matched across experimental groups. Littermate controls were used when possible.

The following mouse strains were obtained from The Jackson Laboratory: C57BL/6J (stock no. 000664; WT), B6.129P2-Gt(ROSA)26Sortm1(DTA)Lky/J (stock no. 009669; *R26-DTA*), C57BL/6-Gt(ROSA)26Sortm1(HBEGF)Awai/J(stock no. 007900; *R26-iDTR*),B6.Cg-Gt(ROSA)26Sortm9(CAG-tdTomato)Hze/J (stock no. 007909; *R26-tdTomato*), B6.Cg-Tg(tetO-Cre)1Jaw/J (stock no. 006234; tetO-Cre), B6.Cg-Gt(ROSA)26Sortm6(CAG-ZsGreen1)Hze/J (stock no. 007906; *R26-ZsGreen*), B6.129-Trpv1tm1(cre)Bbm/J (stock no. 017769; *Trpv1-Cre*), B6N;129-Tg(CAG-CHRM3*,-mCitrine)1Ute/J (stock no. 026220; *R26-hM3Dq*), B6J.Cg-Gt(ROSA)26Sortm96(CAG-GCaMP6s)Hze/MwarJ (stock no. 028866; *R26-GCaMP6s*), B6.129S7-Rag1tm1Mom/J (stock no. 002216; *Rag1*−/−), C;129S4-Rag2tm1.1Flv Il2rgtm1.1Flv/J (stock no. 014593; *Rag2^−/−^Il2rg^−/−^*), B6.129P2-Cxcr4tm2Yzo/J (stock no. 008767; *Cxcr4^flox^*), and B6;129S7-Wnt5atm1Amc/J (stock no. 004758; *Wnt5a^+/−^*). *Ramp1^flox/flox^*mice were provided by Dr. Isaac Chiu (Harvard Medical School), *Iqgap3-CreERT2* mice by Dr. Yoshiaki Ito (National University of Singapore)^20^, B6(C)-Il5tm1.1(icre)Lky/J (*Red5)* mice by Dr. Richard Locksley (University of California, San Francisco)^12^, and *LSL-Kras^G12D^*mice by Dr. Kenneth Olive (Columbia University). *Mist1-CreERT2; CDH1^flox/flox^; LSL-RHOA^Y42C^* mice were described previously^21^. All strains were maintained on a C57BL/6 background and intercrossed as indicated.

For induction of CreERT2-mediated recombination, tamoxifen (Sigma) was administered at 100 mg kg⁻¹ unless otherwise specified. This regimen was used for all CreERT2 mouse lines except in experiments involving HDT-induced gastric injury, in which mice received the high-dose tamoxifen protocol described following.

Sample sizes were based on prior experience with these models and were sufficient to detect biologically meaningful differences. Investigators were not blinded during experiments. No statistical methods were used to predetermine sample size.

#### HDT-induced gastric injury and recall model

Age- and sex-matched 8–12-week-old mice were subjected to high-dose tamoxifen (HDT)-induced gastric injury. Tamoxifen (Sigma) was dissolved in corn oil and administered by oral gavage at 300 mg kg⁻¹ once daily for 3 consecutive days. Control mice received vehicles alone.

For recall experiments, mice were allowed to recover for at least 3 weeks after the initial HDT injury (1HDT) before re-challenge with the same regimen (2HDT). Gastric tissues were collected at the indicated time points after primary or recall injury.

#### Helicobacter pylori infection and eradication

Helicobacter pylori *SSI* strain was cultured under microaerophilic conditions and administered by oral gavage at 2 × 10⁹ colony-forming units (CFU) in 0.2 ml Brucella broth (Becton Dickinson, Cat# 211088). Mice received 3 doses at 2-day intervals.

For eradication experiments, mice were infected for 8 weeks before antibiotic treatment. Eradication was performed using clarithromycin (7.15 mg kg⁻¹ day⁻¹; Sigma-Aldrich), metronidazole (14.2 mg kg⁻¹ day⁻¹; Sigma-Aldrich), and omeprazole (400 μmol kg⁻¹ day⁻¹; Sigma-Aldrich). Omeprazole was dissolved in 200 μl of 2.5% hydroxypropyl methylcellulose and administered by oral gavage. Antibiotics were dissolved in 200 μl PBS and administered by oral gavage 45 min after omeprazole. Treatments were given twice daily for 7 consecutive days. Successful eradication was confirmed by quantitative PCR analysis of gastric tissues as described below.

#### Acetic acid-induced gastric ulcer model

For induction of acute gastric ulcers, mice were anesthetized and subjected to midline laparotomy. The gastric corpus was gently externalized, and 100% acetic acid was applied to the serosal surface using a capillary tube for 15 s. The acid was immediately removed with a sterile cotton swab, the area was rinsed with sterile saline, and the stomach was returned to the abdominal cavity. The peritoneum and skin were closed in separate layers. Mice were monitored postoperatively and euthanized at the indicated time points for tissue collection.

#### Cell culture

Mouse gastric cancer cells (ACKP cells), derived from *Atp4b-Cre; Cdh1^fl/fl^; Kras^G12D^; Trp53^fl/fl^; YFP* mice, were kindly provided by Dr. Sandra W. Ryeom (Columbia University)^17^. Cells were cultured in DMEM supplemented with 10% FBS and 1% penicillin–streptomycin at 37 °C in a humidified incubator with 5% CO₂. All cells were routinely tested and confirmed to be negative for Mycoplasma contamination.

#### Orthotopic ACKP gastric tumor model

Mice were anesthetized with 2–4% isoflurane and placed on a temperature-controlled heating pad. After midline laparotomy, the anterior gastric wall was gently exteriorized and ACKP gastric cancer cells (1 × 10⁶ cells in 10 μl PBS) were injected into the subserosal layer using a sterile 31G × 5/8-inch insulin syringe, taking care to avoid leakage or peritoneal dissemination. The injection site was briefly observed to confirm successful delivery. The stomach was returned to the abdominal cavity, and the peritoneum and skin were closed in separate layers. Tumor-bearing mice were monitored regularly and euthanized 2 weeks after implantation unless humane endpoints were reached earlier. Tumor establishment was confirmed by gross and histological analysis.

All tumor growth studies were performed in accordance with IACUC guidelines. Mice were euthanized when tumor volume exceeded 500 mm³ or when predefined clinical endpoints were met. These limits were not exceeded in any experiments.

#### Nodose ganglion transplantation

Donor nodose ganglia (NG) were isolated from age-matched mice under sterile conditions. Briefly, mice were anesthetized, bilateral NGs were dissected under a stereomicroscope, and ganglia were maintained in ice-cold sterile HBSS (Thermo Fisher Scientific) before transplantation.

Recipient mice were anesthetized with 2–4% isoflurane and placed on a temperature-controlled heating pad. After midline laparotomy, a small subserosal pocket was created in the anterior gastric wall using fine forceps. A single donor NG was inserted into the pocket, and 15 μl growth factor-reduced Matrigel (Corning) was applied to secure the graft and facilitate local stabilization and axonal outgrowth. The stomach was returned to the abdominal cavity, and the peritoneum and skin were closed in separate layers. Control mice underwent sham surgery consisting of identical surgical exposure without NG transplantation. Recipient mice were allowed to recover for 4 weeks before subsequent HDT challenge or orthotopic tumor implantation and were euthanized at the specified time points for analysis. Because donor and recipient mice were maintained on a C57BL/6 background, no immunosuppressive treatment was required.

NG graft survival was evaluated by histological examination of the transplantation site. Successful grafts were defined by the presence of an identifiable nodose ganglion structure containing viable neuronal cell bodies and associated tdTomato⁺ axonal projections extending into the surrounding gastric tissue. Grafts that could not be identified or lacked detectable neuronal structures were classified as failed grafts.

#### AAV injection into the left nodose ganglion

For NG-targeted viral delivery, mice were anesthetized with 2–4% isoflurane and placed on a heating pad. A ventral cervical incision was made, and the left NG was exposed under a stereomicroscope. A 33G Hamilton syringe was used to inject AAV vectors directly into the left NG. A total of 500 nl virus (1 × 10¹² vg ml⁻¹) or vehicle (PBS) was delivered slowly to minimize reflux and tissue damage, and the needle was left in place for 1–2 min before withdrawal.

The incision was closed in layers, and mice were allowed to recover for 2–3 weeks before subsequent procedures. Unless otherwise indicated, analyses were performed on the anterior gastric wall innervated by the injected left nodose ganglion. Control animals underwent identical NG injections with the corresponding control AAV vectors (AAV-Control) matched for serotype, promoter, and fluorescent reporter.

All AAVs were packaged as AAV2/9 serotype vectors. The following viral constructs were used: rAAV-cFos-tTA-pA (Addgene), rAAV-hSyn-FLEX-TeLC-P2A-dTomato (Addgene), rAAV-hSyn-FLEX-mGFP-P2A-Synaptophysin-mRuby (Addgene), rAAV-Syn-ChrimsonR-tdTomato (Addgene), rAAV-U6-sgSmyd4-GFP (ABM), rAAV-CAG-mCherry-U6-mIl13ra1-shRNA (VectorBuilder), and rAAV-CAG-mCherry-U6-mIl5ra-shRNA (VectorBuilder).

#### Fos-dependent Tet-off activity labeling and targeted manipulation

To label or manipulate injury-activated sensory neurons in the nodose ganglion (NG), a Fos-dependent Tet-off strategy was used. rAAV-cFos-tTA-pA was injected into the left NG as described above. In *tetO-Cre* mice, neuronal activation during primary HDT injury drove tTA-dependent Cre recombination in the absence of doxycycline (DOX). To restrict recombination to the primary injury window, DOX was administered from day 14 after the first HDT injury (1HDT), thereby preventing labeling of neurons activated during recall (2HDT).

For activity-dependent neuronal ablation, rAAV-cFos-tTA-pA was injected into *tetO-Cre; R26-DTA* mice. DOX was withheld during 1HDT and introduced on day 14 to restrict DTA expression to primary injury–activated neurons. In control mice, DOX was administered from 7 days before 1HDT and maintained throughout.

For activity-dependent gene deletion, rAAV-cFos-tTA-pA and rAAV-U6-sgSmyd4-GFP were co-injected into the NG of *tetO-Cre; LSL-Cas9* mice. DOX was administered from day 14 after 1HDT and maintained thereafter (sgCtrl and sgSMYD4 groups), or from 7 days before 1HDT throughout the experiment (sgSMYD4 control).

Doxycycline (2 mg ml⁻¹) was provided in drinking water supplemented with 5% sucrose and protected from light.

#### Synaptophysin labeling of sensory neuron terminals

To visualize presynaptic structures of sensory neurons innervating the stomach, rAAV-hSyn-FLEX-mGFP-2A-Synaptophysin-mRuby was injected into the left NG as described above. Briefly, 500 nl virus was delivered into the left NG using a 33G Hamilton syringe. Mice were allowed to recover for 3 weeks to permit axonal transport and expression of synaptophysin-mRuby in peripheral nerve terminals.

#### Monosynaptic retrograde rabies tracing

Monosynaptic retrograde tracing was performed using an EnvA-pseudotyped glycoprotein-deleted rabies virus (EnvA-ΔG-RV) system.

To restrict rabies infection to defined gastric epithelial populations, a Cre-dependent helper AAV encoding TVA and optimized rabies glycoprotein (oG) (rAAV-DIO-Ef1a-TVA-2A-oG, 1 × 10¹² vg ml⁻¹, 10 μl per mouse; Addgene) was delivered to the anterior gastric wall of *Iqgap3-CreERT2* mice.

Three days after helper AAV delivery, mice were subjected to HDT-induced gastric injury or LDT. One week after HDT treatment, EnvA-ΔG-dsRed rabies virus (10⁸ IFU ml−1, 10 μl per mouse; Biohippo) was injected into the same gastric region. Mice were euthanized 7 days later. The left NG was dissected and processed for fluorescence imaging to identify retrogradely labeled neurons.

#### Unilateral vagotomy

For unilateral vagotomy, mice were anesthetized with isoflurane and placed on a heated surgical platform. A midline laparotomy was performed to expose the stomach and lower esophagus. The anterior truncal vagus nerve was identified along the esophagus and transected under a stereomicroscope using fine microsurgical scissors, taking care to avoid injury to surrounding vasculature and the posterior vagal trunk. Sham-operated mice underwent identical surgical exposure without nerve transection. Control mice underwent sham surgery consisting of identical surgical exposure without transection of the vagus nerve.

#### Exogenous CGRP administration

To assess the sufficiency of CGRP signaling, recombinant mouse CGRP peptide (Tocris) was administered intraperitoneally at 2 μg per mouse per day beginning on day 1 after HDT injury and continued daily through day 7. Control mice received vehicle injections. Mice were euthanized at the indicated time points for histological and tumor analyses.

#### Chemogenetic activation of Trpv1⁺ neurons

For chemogenetic activation of sensory neurons, *Trpv1-Cre; R26-hM3Dq* mice were administered clozapine-N-oxide (CNO, Tocris) by oral gavage at 0.3 mg kg⁻¹ once daily beginning on day 1 after HDT injury and continued daily through day 7. Control mice received vehicle gavage. Gastric tissues were collected at the indicated time points.

#### CGRP receptor antagonism

To inhibit CGRP signaling during recall or tumor progression, Rimegepant (MedChem Express, HY-15498) was administered intraperitoneally at 4 mg kg⁻¹ once daily beginning on day 1 after the second HDT injury or day 1 after orthotopic ACKP tumor implantation and continued daily for 14 consecutive days. Control mice received vehicle injections.

#### Methionine restriction and betaine supplementation

Methionine restriction (MR) diet was used to modulate systemic methyl donor availability. Mice were switched to a methionine-restricted diet containing 0.12% methionine (Research Diets) at the indicated time points and maintained for the duration specified in each experiment. Control mice received an isocaloric control diet containing 0.86% methionine (Research Diets).

For betaine rescue experiments, betaine (Sigma-Aldrich) was administered in drinking water at 1% (w/v) starting concurrently with the MR diet and maintained throughout the experimental period. In HDT-induced injury experiments, mice were placed on the MR diet before primary injury and maintained through the recall phase as indicated.

In *Helicobacter pylori* eradication experiments, MR diet was initiated during antibiotic treatment and continued for 2 weeks. Body weight and general health status were monitored throughout dietary intervention.

#### Magnetic resonance imaging (MRI)

Magnetic resonance imaging (MRI) was performed on a Bruker 9.4 T preclinical scanner using ParaVision 7 software for image acquisition and analysis. Mice were anesthetized with isoflurane in medical air, and body temperature was maintained throughout imaging using the physiological monitoring system.

MRI scans were acquired at baseline (day 0) before injury and again at the indicated time points after injury. After acquisition of axial T2-weighted anatomical images for localization, T1 mapping and diffusion-weighted imaging (DWI) were performed in the same axial slice locations. Regions of interest (ROIs) corresponding to the NTS were manually delineated on axial images based on anatomical landmarks, and T1, T2 and ADC values were extracted for quantitative analysis using ParaVision 7 software.

#### In vivo optogenetic stimulation of NG neurons coupled with gastric Iqgap3⁺ epithelial Ca²⁺ imaging

In vivo calcium imaging of gastric epithelial cells during vagal sensory neuron activation was performed using *Iqgap3-CreERT2; R26-GCaMP6s; R26-tdTomato* mice. To enable optogenetic activation of vagal afferents, the left NG was injected with rAAV-Syn-ChrimsonR-tdTomato as described above, and experiments were performed 3 weeks later to allow robust expression. To label Iqgap3⁺ gastric progenitor cells, mice received either low-dose tamoxifen (LDT) or high-dose tamoxifen (HDT) 2 days before calcium imaging experiments.

Mice were anesthetized with isoflurane and placed supine on a heated stage. The left NG was exposed by blunt dissection. For epithelial imaging, the abdominal cavity was opened, and the anterior gastric wall was exteriorized and stabilized for real-time microscopy. Optogenetic stimulation was delivered to the exposed left NG using red light through a fiber-coupled light source (630–635 nm). During NG photostimulation, time-lapse imaging of GCaMP6s fluorescence in Iqgap3-lineage (tdTomato⁺) gastric epithelial cells was acquired in vivo. GCaMP6s signals were recorded with 488-nm excitation and appropriate emission collection, and tdTomato fluorescence was used to identify Iqgap3-lineage cells.

#### Calcium imaging of NG neurons (ex vivo)

Calcium imaging was performed on freshly isolated nodose ganglia (NG) from *Trpv1-Cre; R26-GCaMP6s* mice. NGs were immersed in Live Cell Imaging Solution (Thermo Fisher) and maintained at 37 °C during imaging. Time-lapse imaging was performed using a Leica TCS SP8 upright confocal microscope in XYZT acquisition mode at a temporal resolution of 3 s and a spatial resolution of 512 × 512 pixels. GCaMP6s fluorescence was excited at 488 nm and emission collected at 500–550 nm.

After establishing a stable baseline, stimuli were applied by bath addition as indicated, including recombinant mouse IL-5 (10 ng ml⁻¹, PeproTech), IL-13 (10 ng ml⁻¹, PeproTech), amphiregulin/AREG (50 ng ml⁻¹, R&D Systems), Wnt5a (200 ng ml⁻¹, R&D Systems), and capsaicin (1 µM, Sigma-Aldrich). Regions of interest corresponding to individual Trpv1⁺ neurons were selected, and responses were quantified as ΔF/F₀ relative to the pre-stimulus baseline. Neurons were defined as responsive when ΔF/F₀ exceeded baseline fluctuations, and the fraction of responsive neurons and response amplitude were quantified for each condition.

#### Flow cytometry and cell sorting

Stomach or gastric tumor tissues were collected and flushed with cold PBS, then minced into small fragments. Tissues were digested in digestion medium (HBSS supplemented with 1% HEPES, 1 mg ml⁻¹ Collagenase P, 1 mg ml⁻¹ Dispase II, 10 μg ml⁻¹ DNase I and 1% BSA) at 37 °C for 20 min with gentle agitation. The resulting cell suspension was filtered through a 40 μm strainer and subjected to red blood cell lysis using RBC lysis buffer (BioLegend) for 5 min. The reaction was quenched with PBS. Cells were then stained with TruStain FcX (BioLegend) for 5 min, followed by incubation with primary antibodies for 30 min and fluorophore-conjugated secondary antibodies where applicable for 30 min (1:200).

For flow cytometric analysis, cells were first gated on singlets and live cells using Zombie NIR viability dye (BioLegend, 423105). Leukocytes were identified as CD45⁺ cells (BV421 anti-mouse CD45, BioLegend, 103134).

Major immune populations were defined as follows: T cells, CD45⁺ CD3⁺ (PE/Cy5 anti-mouse CD3, BioLegend, 128617); dendritic cells, CD45⁺ CD11c⁺ (PE anti-mouse CD11c, BioLegend, 117307); macrophages, CD45⁺ CD11b⁺ F4/80⁺ (BV650 anti-mouse CD11b, BioLegend, 101239; APC anti-mouse F4/80, BioLegend, 123116); neutrophils, CD45⁺ CD11b⁺ Ly6G⁺ (PE/Cy7 anti-mouse Ly6G, BioLegend, 127618); monocytes, CD45⁺ CD11b⁺ Ly6C⁺ (BV711 anti-mouse Ly6C, BioLegend, 128037); and innate lymphocytes, CD45⁺ Lineage⁻ CD127⁺ CD90.2⁺ (Alexa Fluor 700 anti-mouse CD45, BioLegend, 103128; Pacific Blue anti-mouse Lineage Cocktail, BioLegend, 133305; BUV737 anti-mouse CD127, BD Biosciences, 612841; BV605 anti-mouse CD90.2, BioLegend, 105343). The lineage cocktail included antibodies against CD3 (clone 17A2), Ly6G/Ly6C (clone RB6-8C5), CD11b (clone M1/70), CD45R/B220 (clone RA3-6B2), and TER-119 (clone TER-119).

ILC2 populations were identified as CD45⁺ Lineage⁻ CD127⁺ CD90.2⁺ SCA1⁺ ICOS⁺ cells (BV421 anti-mouse Ly6A/E, BD Biosciences, 741222; BV785 anti-mouse CD278, BioLegend, 313533). Gastric ILC2 subsets were further distinguished based on IL-33R (BUV395 anti-mouse ST2, BD Biosciences, 745738) and IL-25R (BV480 anti-mouse IL-17 Receptor B, BD Biosciences, 746606) expression as indicated.

For intracellular cytokine staining, cells were stimulated for 6 h using BD Leukocyte Activation Cocktail with BD GolgiPlug (BD Biosciences, 550583), fixed and permeabilized using the BD Cytofix/Cytoperm Fixation/Permeabilization Solution Kit (BD Biosciences, 554714), and stained for intracellular IL-5 (APC anti-mouse IL-5, BioLegend, 504306) and IL-13 (PE anti-mouse IL-13, BD Biosciences, 568551).

For ILC2 sorting experiments, gastric ILC2s were isolated from Red5 reporter mice, in which IL-5–producing ILC2s are labeled by tdTomato. Briefly, gastric tissues were enzymatically digested to generate single-cell suspensions, and ILC2s were sorted based on CD45⁺ (Alexa Fluor 700 anti-mouse CD45, BioLegend, 103128), Lineage⁻ (Pacific Blue anti-mouse Lineage Cocktail, BioLegend, 133305), CD127⁺ (FITC anti-mouse CD127, BioLegend, 135007), CD90.2⁺ (BV605 anti-mouse CD90.2, BioLegend, 105343), SCA1⁺ (PerCP-Cy5.5 anti-mouse Ly6A/E, BioLegend, 122523), and ICOS⁺ (BV785 anti-mouse CD278, BioLegend, 313533) surface markers.

For epithelial cell sorting, gastric epithelial cells were isolated from *Iqgap3-CreERT2; R26-tdTomato* mice. Dead cells were excluded using DAPI, and epithelial lineage cells were identified as EpCAM⁺ (APC anti-mouse CD326/EpCAM, BioLegend, 118214) tdTomato⁺ cells prior to sorting.

All cell sorting experiments were performed using a BD Influx cell sorter. Flow cytometry data acquisition was performed on a NovoCyte Quanteon or NovoCyte Penteon flow cytometer, and data were analyzed using FlowJo software (v.10).

#### ILC2 culture

For in vitro expansion of ILC2s, gastric ILC2s were purified by fluorescence-activated cell sorting (FACS) as described above. Sorted ILC2s were seeded at 500 cells per well in 200 μl RPMI-1640 medium (Thermo Scientific) supplemented with 1× MEM non-essential amino acids (Thermo Scientific), 1 mM sodium pyruvate, 100 IU ml⁻¹ penicillin–streptomycin (Thermo Scientific), 10 mM HEPES (Cytiva), 1× 2-mercaptoethanol (Thermo Scientific), 10% FBS (Sigma-Aldrich), 10 ng ml⁻¹ IL-2 (PeproTech), and 10 ng ml⁻¹ IL-7 (BioLegend). Cells were cultured at 37 °C in a humidified incubator with 5% CO₂ and used for downstream coculture experiments or generation of ILC2-conditioned medium as indicated.

#### Gastric epithelial organoid culture

For gastric epithelial organoid cultures, epithelial cells were isolated from gastric tissues and purified by FACS as described above. EpCAM⁺ tdTomato⁺ cells from *Iqgap3-CreERT2; R26-tdTomato* mice were collected and used for organoid culture.

A total of 2,000 sorted single epithelial cells were mixed with 20 μl growth factor-reduced Matrigel (Corning) per well and plated in 24-well plates. After Matrigel polymerization at 37 °C, organoid cultures were maintained in basal medium consisting of 50% conditioned medium containing Wnt3A, R-spondin-1, and Noggin (WRN, ATCC) and 50% Advanced DMEM/F12 (Gibco) supplemented with 20% FBS (Sigma-Aldrich), B27 (1×, Gibco), N2 (1×, Gibco), and 1% penicillin–streptomycin (Thermo Scientific). Medium was additionally supplemented with EGF (50 ng ml⁻¹, Gibco), FGF10 (100 ng ml⁻¹, PeproTech), Y-27632 (10 μM, Sigma-Aldrich), GlutaMAX (2 mM, Gibco), HEPES (10 mM), and SB431542 (10 μM, Tocris). Medium was replaced every other day. Organoids were passaged every 6 days at a 1:3 ratio by dissolving Matrigel with TrypLE (Gibco) followed by mechanical dissociation by pipetting.

#### Nodose ganglion culture and neurite outgrowth assay

Nodose ganglia (NG) were freshly isolated from adult mice (8–12 weeks old) and embedded in growth factor-reduced Matrigel (Corning) as 20 μl droplets in prewarmed 24-well plates. After polymerization for 20 min at 37 °C, wells were overlaid with culture medium. Recombinant mouse AREG (50 ng ml⁻¹), NGF (50 ng ml⁻¹), or Wnt5a (100 ng ml⁻¹) was added as indicated. Control wells received vehicle treatment. NGs were cultured for 3–5 days, then fixed and stained for TUBB3 to visualize neuronal processes. Axonal outgrowth was quantified using an outgrowth index calculated as the ratio of the axon-positive area fraction in the distal outgrowth region to that in the soma-adjacent region.

#### Nodose ganglion–organoid coculture

Gastric organoids were embedded in Matrigel domes as described above and allowed to solidify. Freshly isolated NGs were positioned adjacent to the organoid-containing Matrigel dome (approximately 0.5–1 mm from the dome edge) and embedded in an additional Matrigel droplet. Cocultures were maintained in organoid growth medium, and organoid diameter and neuron–organoid contact numbers were quantified on day 5 unless otherwise indicated.

#### Nodose ganglion–ILC2–organoid triple coculture

For triple coculture assays, gastric organoids were first embedded in Matrigel domes in the lower chamber of a transwell system. Freshly isolated NGs were positioned adjacent to the organoid dome and embedded in Matrigel in the lower chamber. IL-25R⁺ gastric ILC2s were sorted by flow cytometry and seeded into the upper transwell insert (5 × 10⁴ cells per well), allowing paracrine signaling without direct cell–cell contact. In selected experiments, ILC2s were isolated from naïve mice or from mice subjected to HDT-induced injury. Cocultures were maintained for up to 5 days, and organoid diameter, neurite extension, and neuron–organoid contact formation were quantified by confocal microscopy.

#### ILC2 depletion in *Red5; iDTR* mice

To selectively ablate IL-5–expressing ILC2s, Red5; iDTR mice were administered diphtheria toxin (DT; EMD Millipore) intraperitoneally at 250 ng every other day for a total of three injections. For ILC2 depletion during the establishment phase of inflammatory memory, DT administration was initiated 10 days before the first HDT injury. For depletion during the maintenance phase, DT administration was initiated 10 days after completion of the first HDT injury. Unless otherwise indicated, HDT-induced injury and subsequent procedures were performed according to the depletion schedule. Efficient depletion of ILC2s was confirmed by flow cytometric analysis of gastric CD45⁺ Lineage⁻ CD127⁺ CD90.2⁺ SCA1⁺ ICOS⁺ cells as described above.

#### Adoptive transfer of ILC2s

For ILC2 adoptive transfer experiments, gastric ILC2s were isolated from Red5 reporter donor mice. Briefly, gastric tissues were enzymatically digested to generate single-cell suspensions, and ILC2s were sorted based on CD45⁺ (Alexa Fluor 700 anti-mouse CD45, BioLegend, 103128), Lineage⁻ (Pacific Blue anti-mouse Lineage Cocktail, BioLegend, 133305), CD127⁺ (FITC anti-mouse CD127, BioLegend, 135007), CD90.2⁺ (BV605 anti-mouse CD90.2, BioLegend, 105343), SCA1⁺ (PerCP-Cy5.5 anti-mouse Ly6A/E, BioLegend, 122523), and ICOS⁺ (BV785 anti-mouse CD278, BioLegend, 313533) surface markers.

A total of 1 × 10⁵ ILC2s were injected intravenously into each mouse, with 3 injections given within 1 week (3 × 10⁵ total ILC2s per mouse). Recipient *Rag2^−/−^Il2rg^−/−^* mice were subjected to HDT injury or orthotopic tumor implantation as indicated. Engraftment of transferred ILC2s in recipient gastric tissues was confirmed by detection of tdTomato⁺ cells by immunofluorescence microscopy at the indicated time points.

#### Immune cell depletion experiments

To assess the contribution of specific immune cell populations to the establishment of inflammatory memory, selective immune cell depletion was performed before the initial HDT injury. Macrophages were depleted by intraperitoneal injection of clodronate liposomes (Encapsula NanoSciences, 200 μl per mouse) 48 h before the first HDT injury; control mice received PBS-loaded liposomes. Neutrophils were depleted using anti-Ly6G monoclonal antibody (BP0075-1, Bio X Cell, 200 μg per mouse, intraperitoneally) 24 h before the first HDT injury; control mice received isotype-matched IgG. Dendritic cells were targeted using anti-CD11c monoclonal antibody (BE0038, Bio X Cell, 200 μg per mouse, intraperitoneally) 24 h h before the first HDT injury; control mice received isotype-matched IgG. Innate lymphoid cells were depleted using anti-CD90.2 monoclonal antibody (BP0066, Bio X Cell, 200 μg per mouse, intraperitoneally) 24 h before the first HDT injury; control mice received isotype-matched IgG. Efficient depletion of target populations was confirmed by flow cytometric analysis of gastric CD45⁺ immune cells at the time of primary injury.

#### Cytokine neutralization

For cytokine neutralization experiments, mice were treated with neutralizing antibodies against IL-5 or IL-13. Anti-IL-5 antibody (Bio X Cell, BE0198) or anti-IL-13 antibody (eBioscience, eBio1316H) was administered by intraperitoneal injection at 200 μg per mouse. Antibodies were administered starting 1 day before injury and continued every 3 days throughout the experimental period as indicated. Control mice received equivalent doses of isotype-matched control IgG.

#### Immunofluorescence

Tissues were fixed overnight in 4% paraformaldehyde or 10% neutral-buffered formalin at 4 °C and embedded in paraffin or OCT compound. Tissue sections were prepared using standard histological procedures. For immunostaining, sections underwent antigen retrieval, were blocked with 10% BSA for 1 h at room temperature, and incubated with primary antibodies overnight at 4 °C.

Primary antibodies used were anti-Trpv1 (1:200, Alomone Labs, ACC-030), anti-CGRP (1:200, Abcam, ab81887 or ab36001), anti-TUBB3 (1:200, Abcam, ab18207), anti-TH (1:200, Sigma-Aldrich, AB152), anti-ChAT (1:200, Sigma-Aldrich, AB144P), anti-Ki67 (1:200, Fisher Scientific, 550609), anti-ATP4B (1:200, Santa Cruz Biotechnology, sc-374094), anti-GIF (1:200, MyBioSource, MBS2028736), GS-II lectin (1:200, Invitrogen, L21415), anti-Ramp1 (1:200, Abcam, ab156575), anti-GATA3 (1:200, Santa Cruz Biotechnology, sc-268), anti-IL25R (1:200, BD Biosciences, 746606), anti-SMYD4 (1:200, Proteintech, 17594-1-AP), anti-H3K4me3 (1:200, Rockland, 600-401-I59), anti-H3K4me1 (1:200, Abcam, ab8895), anti-H3K27ac (1:200, Abcam, ab4729), anti-TROP2 (1:200, R&D Systems, AF1122), anti-Iqgap3 (1:200, Thermo Fisher Scientific, PA5-101817), anti-YFP (1:200, Biorbyt, orb334991), anti-GFP (1:200, Aves Labs, GFP-1010), anti-tdTomato (1:200, Biorbyt, orb182397), anti-mRuby (1:200, Biorbyt, orb1463286), anti-RFP (1:200, Rockland, 600-401-379), anti-IL5RA (1:200, Bioss Antibodies, BS-2601R), anti-IL13RA1 (1:200, Santa Cruz Biotechnology, sc-398831), anti-IL4RA (1:200, Santa Cruz Biotechnology, sc-28361), anti-EGFR (1:200, Abcam, ab52894), anti-FZD3 (1:200, Abcam, ab217032), anti-cFOS (1:200, Cell Signaling Technology, 2250S), anti-phospho-ERK1/2 (pERK) (1:200, Cell Signaling Technology, 4370S), and anti-CD45 (1:200, Abcam, ab10558), as indicated in individual experiments.

After washing, sections were incubated with Alexa Fluor-conjugated secondary antibodies (Alexa Fluor 488, 594 or 647; Thermo Fisher Scientific) for 1 h at room temperature. Nuclei were counterstained with DAPI.

#### RNAscope in situ hybridization

RNAscope in situ hybridization was performed using the RNAscope Multiplex Fluorescent Reagent Kit v2 (Advanced Cell Diagnostics) according to the manufacturer’s instructions. Paraffin-embedded gastric tissue sections were baked in a dry oven for 1 h at 60 °C followed by deparaffinization. Sections were then treated with hydrogen peroxide to block endogenous peroxidase activity. After hydrophobic barriers were drawn around the tissue sections, protease reagent was applied for target retrieval.

RNAscope probes targeting mouse Ramp1 and Iqgap3 transcripts (Advanced Cell Diagnostics, 532681 and 539811) were applied to the sections and incubated in a humidified chamber at 40 °C. After probe hybridization, slides were washed twice in RNAscope wash buffer and sequential signal amplification was performed according to the manufacturer’s protocol. Ramp1 signals were detected using Opal 520 (green channel), whereas Iqgap3 signals were detected using Opal 570 (red channel). Slides were counterstained with DAPI and mounted with antifade mounting medium.

#### Whole-mount tissue clearing and 3D imaging (iDISCO)

Three-dimensional visualization of gastric innervation was performed using the iDISCO tissue clearing method. Mouse stomach tissues were harvested and fixed overnight in 4% paraformaldehyde at 4 °C, dehydrated through a graded methanol series (20%, 40%, 60%, 80%, and 100%), and bleached overnight in methanol containing 5% hydrogen peroxide at 4 °C to reduce tissue autofluorescence.

After rehydration, tissues were permeabilized in PBS containing Triton X-100, DMSO, and glycine, followed by blocking in PBS supplemented with serum and detergent. Samples were incubated with primary antibodies for 3–5 days at 37 °C with gentle agitation. Primary antibodies included anti-tdTomato (Biorbyt, orb182397) and anti-CGRP (Abcam, ab81887) to visualize Iqgap3⁺ cells and sensory nerve fibers. Following extensive washing, tissues were incubated with species-appropriate fluorescent secondary antibodies for 2–3 days at 37 °C, dehydrated again through a graded methanol series, and cleared using dichloromethane and dibenzyl ether. Images were obtained using an Ultramicroscope II light-sheet microscope (2×/0.5 NA objective, 1× zoom) and analyzed using Imaris software according to the manufacturer’s instructions.

#### ELISA assays

For CGRP quantification, nodose ganglia (NG) were cultured ex vivo, and culture supernatants were collected at the indicated time points. Calcitonin gene-related peptide (CGRP) levels were measured using a Mouse CGRP ELISA Kit (MyBioSource, MBS267121) according to the manufacturer’s instructions. Absorbance was measured using a microplate reader, and CGRP concentrations were calculated from standard curves.

For measurement of S-adenosylmethionine (SAM), mouse plasma samples were collected and processed according to the manufacturer’s protocol. SAM levels were quantified using a Mouse S-Adenosylmethionine (SAM) ELISA Kit (MyBioSource, MBS2603023). ELISA signals were measured using a microplate reader and calculated according to standard curves.

#### Neuronal ChIP–qPCR

Chromatin immunoprecipitation followed by quantitative PCR (ChIP–qPCR) was performed using sorted Trpv1⁺ sensory neurons isolated from nodose ganglia (NG).

NGs were dissected from *Trpv1-Cre; R26-tdTomato* mice and enzymatically dissociated. Briefly, NG tissues were incubated at 37 °C in 2 ml HEPES-buffered saline containing collagenase IV (1 mg ml⁻¹, Sigma) and dispase II (2.4 U ml⁻¹, Roche Applied Sciences) for 60 min. The supernatant was removed and replaced with fresh collagenase IV/dispase II solution, followed by a second incubation at 37 °C for 20 min. Cells were transferred to tubes containing 10 ml DMEM supplemented with 10% FBS (Thermo Fisher Scientific) and centrifuged at 200 g for 1 min at 4 °C. The pellet was resuspended in 800 μl DMEM/10% FBS containing DNase I (150 U ml⁻¹, Thermo Fisher Scientific). Cells were gently dissociated into single-cell suspensions using fire-polished glass Pasteur pipettes with progressively reduced tip diameters (VWR International).

tdTomato⁺ sensory neurons were purified by fluorescence-activated cell sorting from *Trpv1-Cre; R26-tdTomato* mice. Typically, 5–10 × 10⁴ sorted Trpv1⁺ neurons were used per ChIP reaction. Chromatin immunoprecipitation was performed using the Magna ChIP™ G – Chromatin Immunoprecipitation Kit (Millipore) according to the manufacturer’s instructions. Sorted neurons were crosslinked with formaldehyde, lysed, and chromatin was sheared by sonication. Immunoprecipitation was performed using antibodies against H3K4me3 (Abcam, AB213224) and H3K27ac (Abcam, ab4729), with normal IgG used as a negative control. Following reverse crosslinking and DNA purification, enriched chromatin fragments were analyzed by quantitative PCR using gene-specific primers. ChIP signals were normalized to input DNA and expressed as fold enrichment relative to IgG controls.

#### Quantitative real-time PCR

Total RNA was extracted from tissues or cultured cells using TRIzol reagent (Invitrogen) according to the manufacturer’s instructions. RNA was reverse transcribed into cDNA using qScript cDNA SuperMix (QuantaBio). Quantitative real-time PCR (qPCR) was performed using SYBR Green FastMix Reaction Mix (QuantaBio) on a 7500 Real-Time PCR System (Applied Biosystems). Relative gene expression levels were calculated using the 2⁻ΔΔCt method and normalized to Gapdh expression. Predesigned primer sequences from Integrated DNA Technologies (IDT) were used for all assays.

#### Statistical analysis and data visualization

Statistical analyses were performed using GraphPad Prism (v.10). Data are presented as mean ± s.e.m., unless otherwise indicated. Quantifications of cell numbers and nerve densities were performed using the average values from each mouse as the biological replicate. Statistical significance between two groups was determined using unpaired two-tailed Student’s t-tests. For multiple-group comparisons, one-way or two-way ANOVA followed by appropriate post hoc tests was used as indicated in the figure legends. Differences in categorical variables, including invasion levels, were analyzed using Fisher’s exact test or the chi-square test as appropriate. Survival curves were analyzed using the Kaplan–Meier method and compared using the log-rank test.

Calcium imaging data were analyzed using Fiji (ImageJ; NIH). H&E-stained sections were analyzed using QuPath (v0.5.1) to delineate lesion areas and quantify the proportion of affected tissue. Public single-cell RNA-seq datasets were analyzed and visualized using the associated online platform. Bulk RNA-seq data were analyzed and visualized using R (v4.2.1). Statistical details are provided in the figures and corresponding legends.

Illustrations were generated with BioRender.com and further edited in Adobe Illustrator 2024.

No statistical methods were used to predetermine sample size, but sample sizes were similar to those generally used in the field. Animals were randomly assigned to experimental groups when possible. Investigators were not blinded during experiments. A *P* value < 0.05 was considered statistically significant. n.s., not significant; \**P* < 0.05, \*\**P* < 0.01, \*\*\**P* < 0.001, \*\*\*\**P* < 0.0001.

---

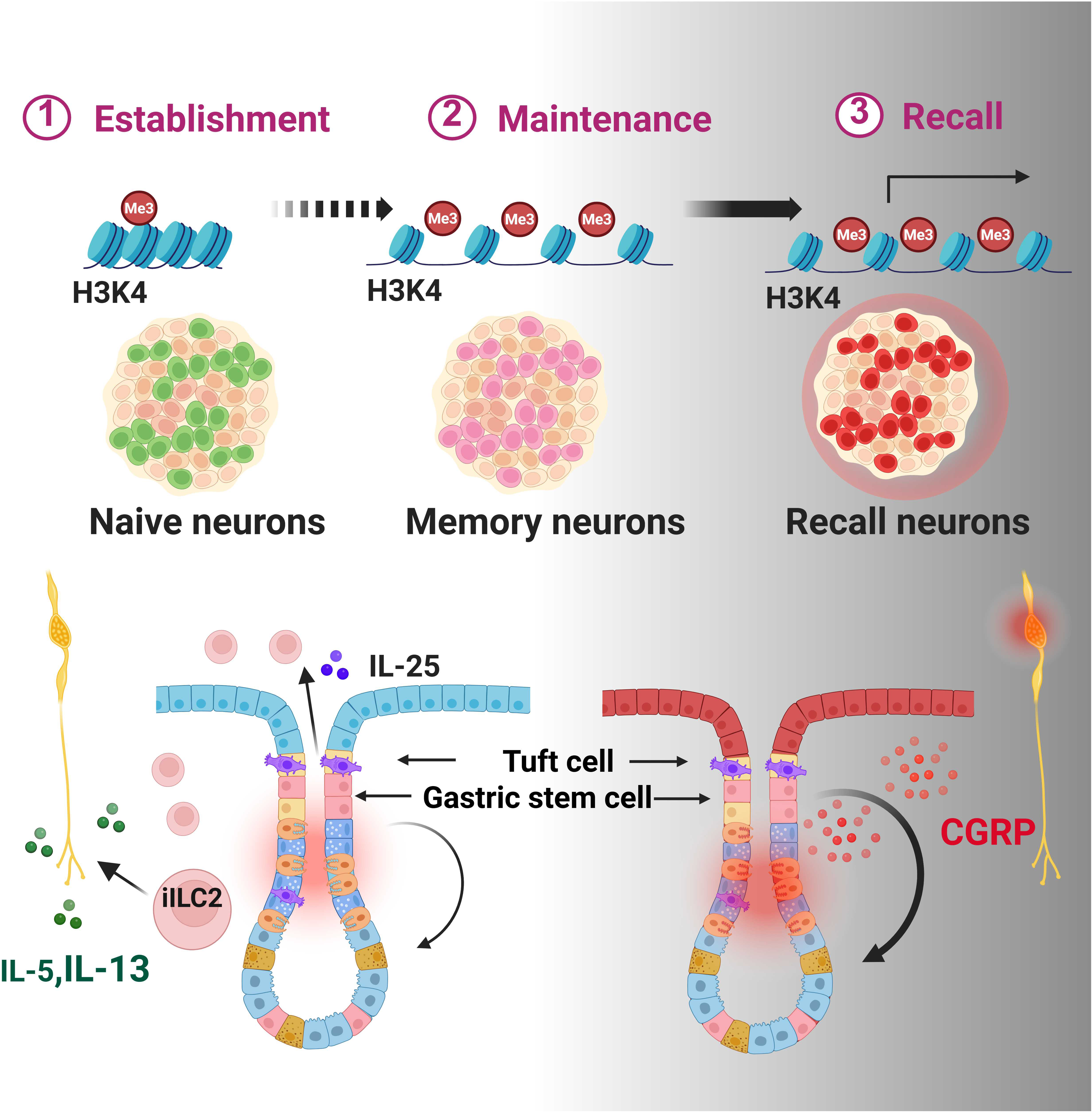

