## Supplementary figures and images for "Sensory neurons encode long-term inflammatory memory that promotes gastric regeneration and tumorigenesis"

### Supplemental figures

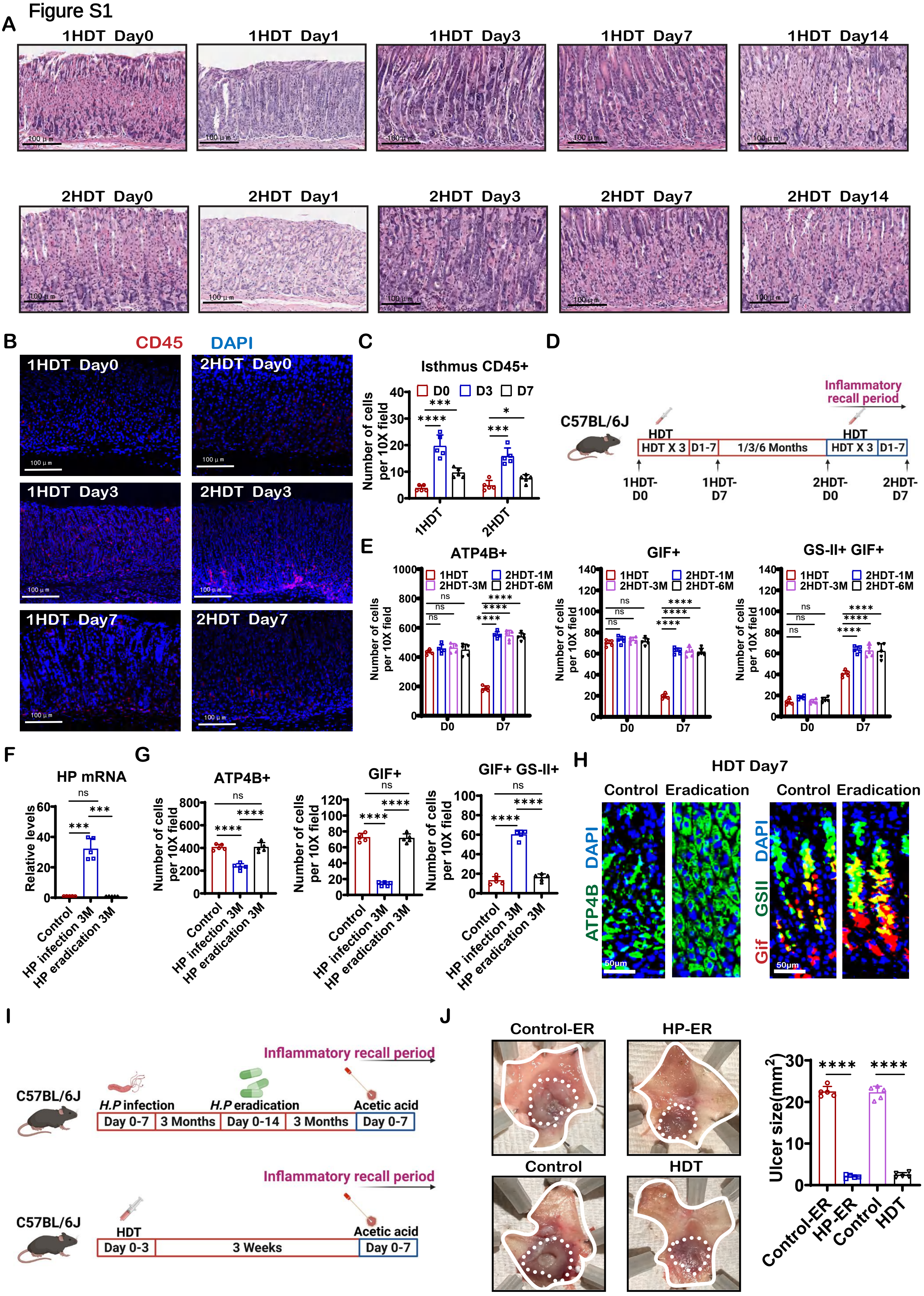

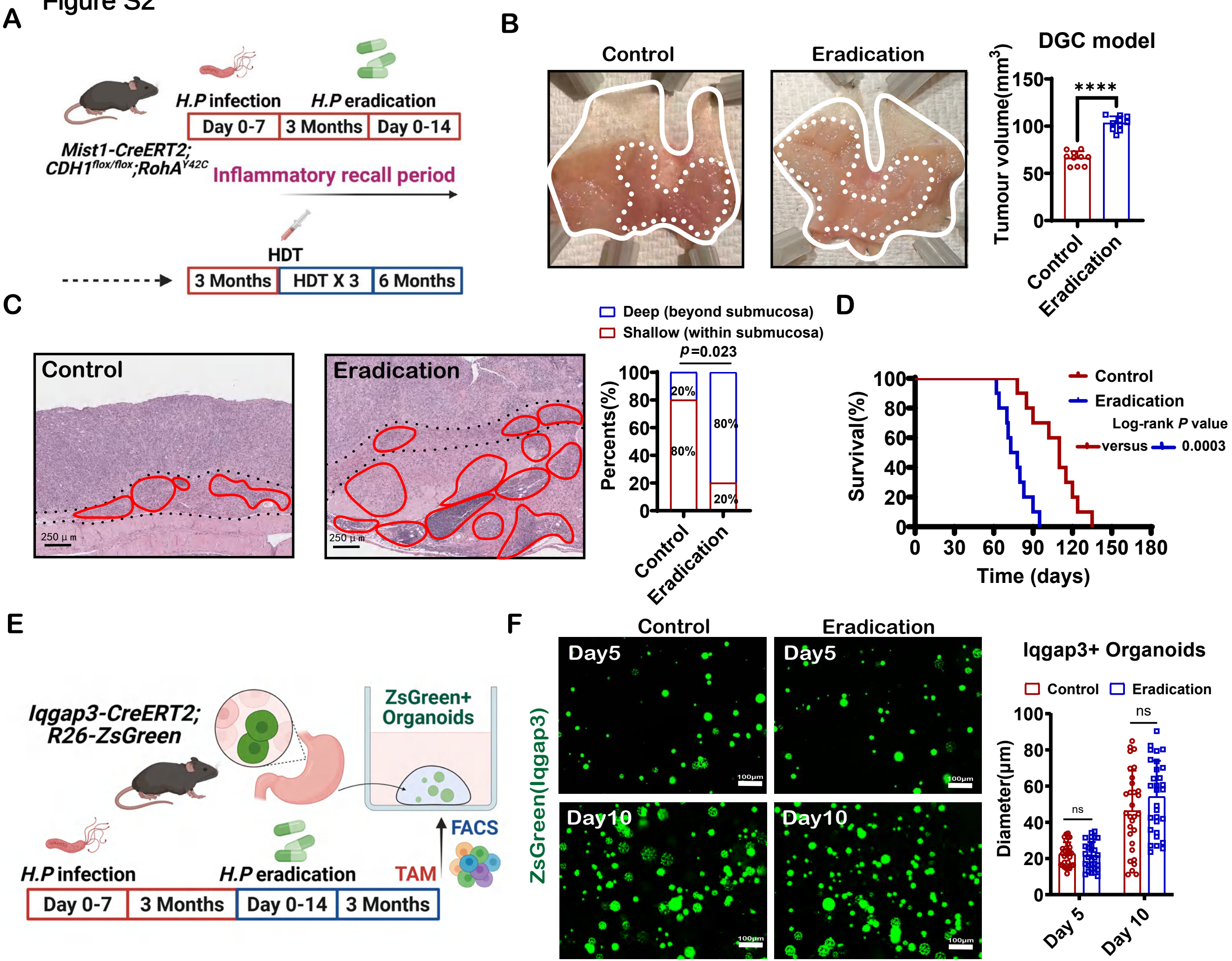

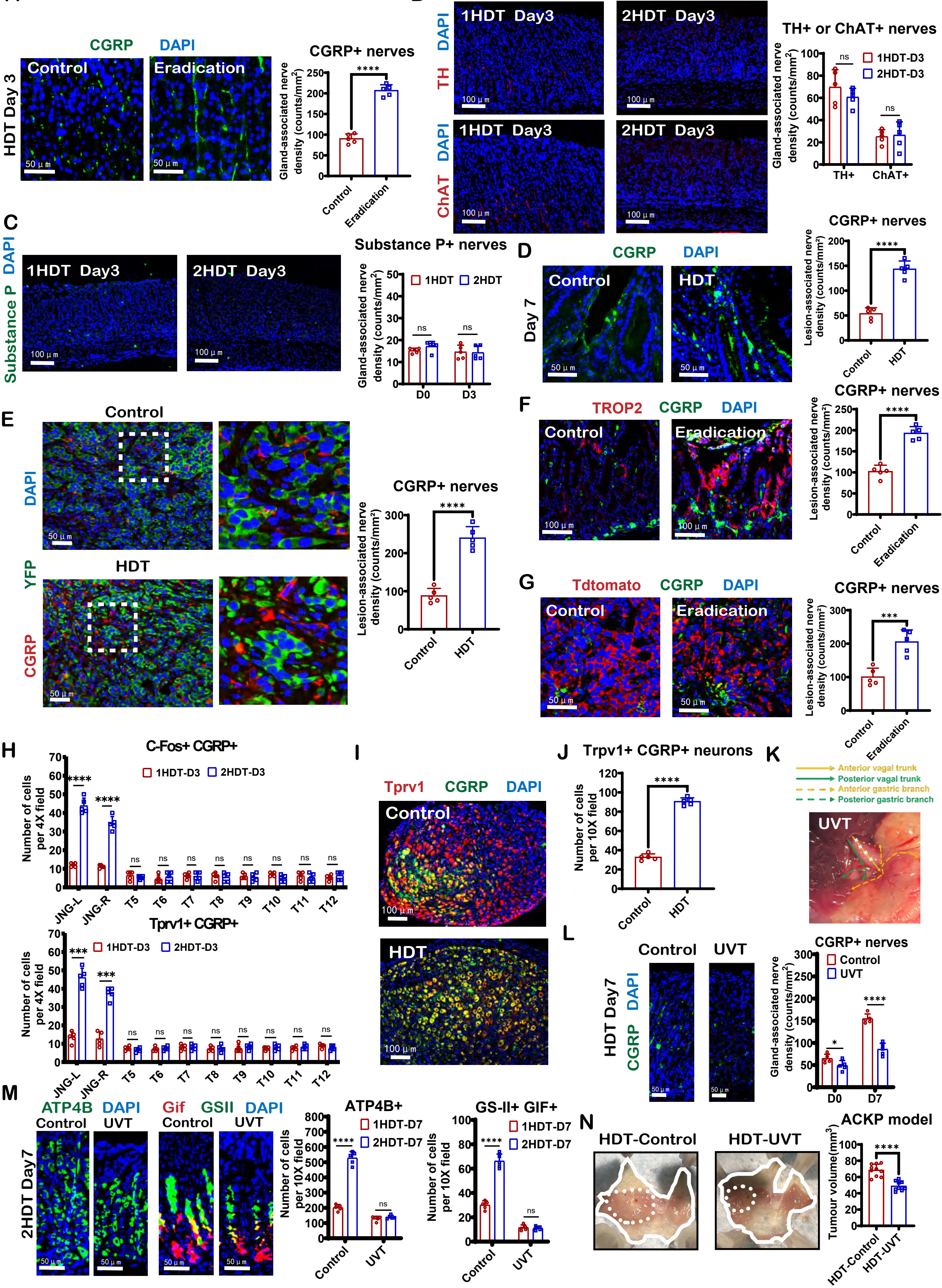

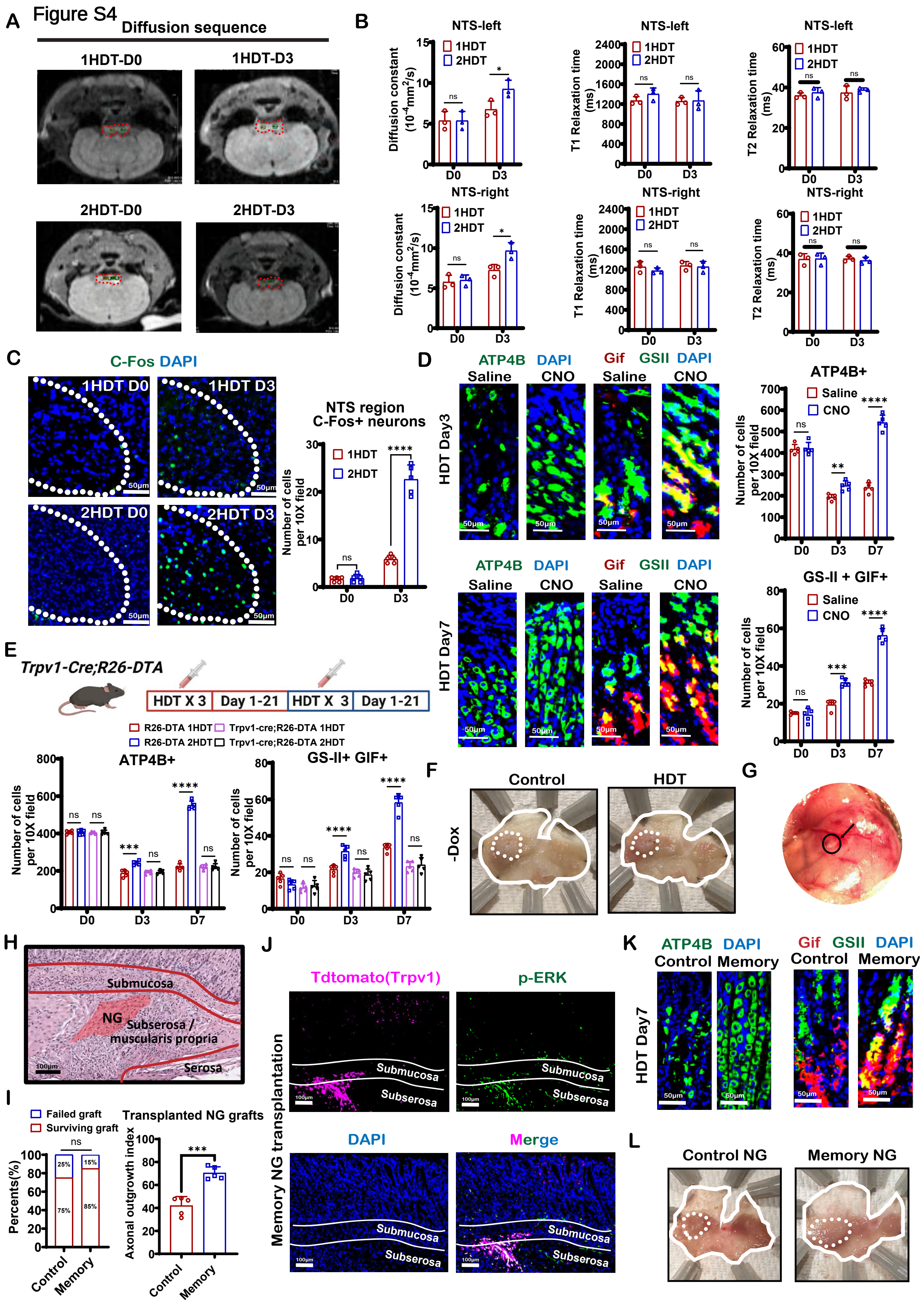

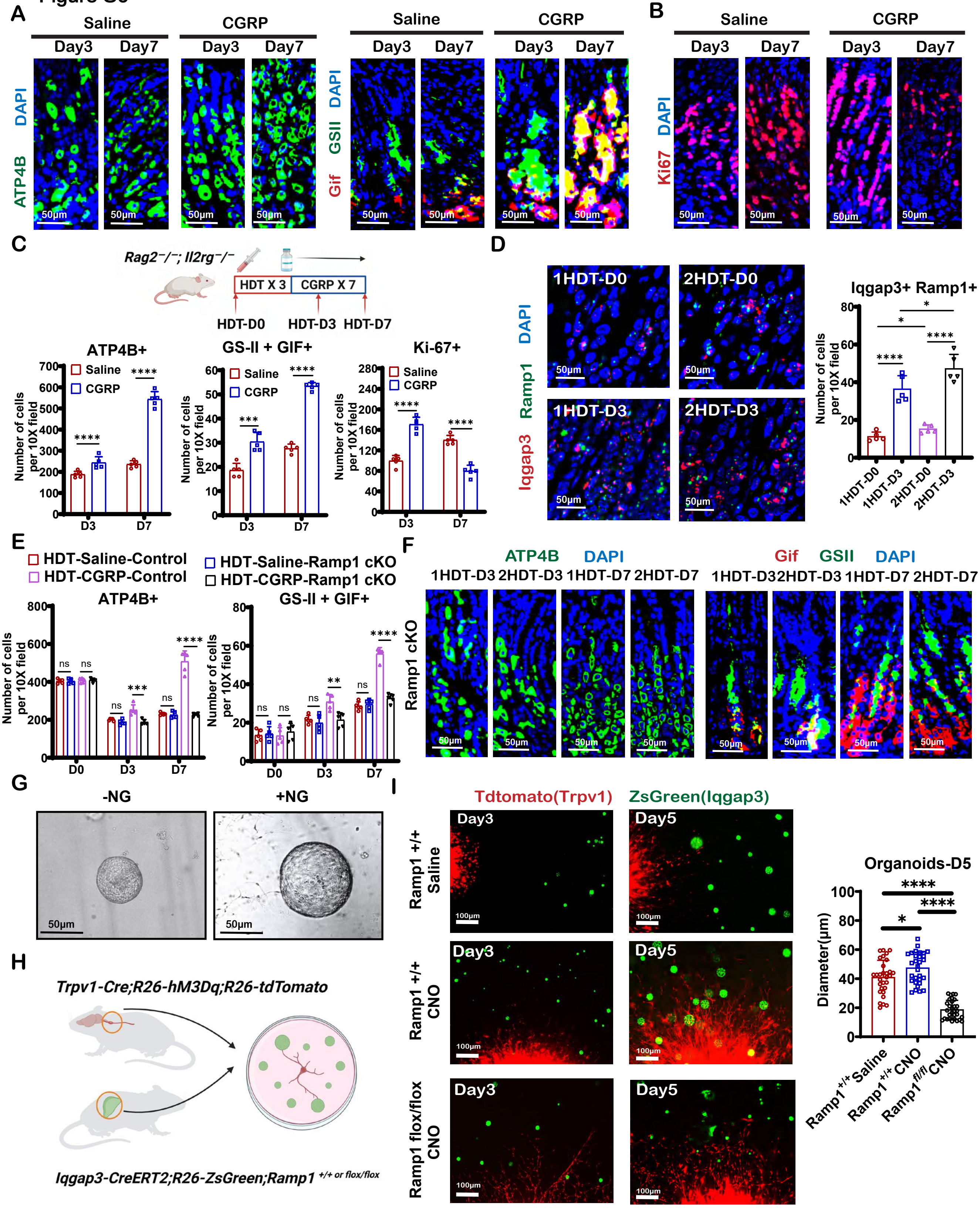

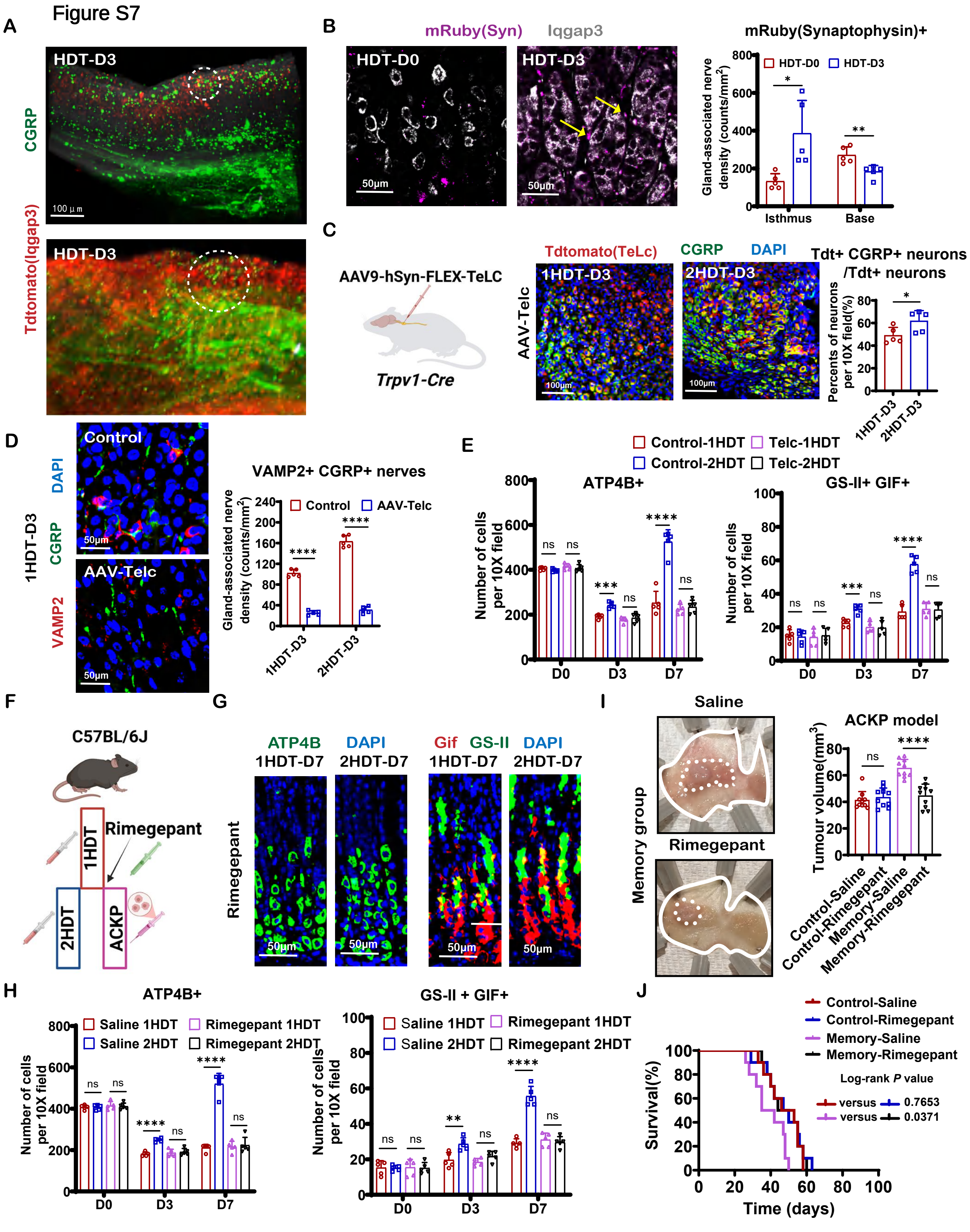

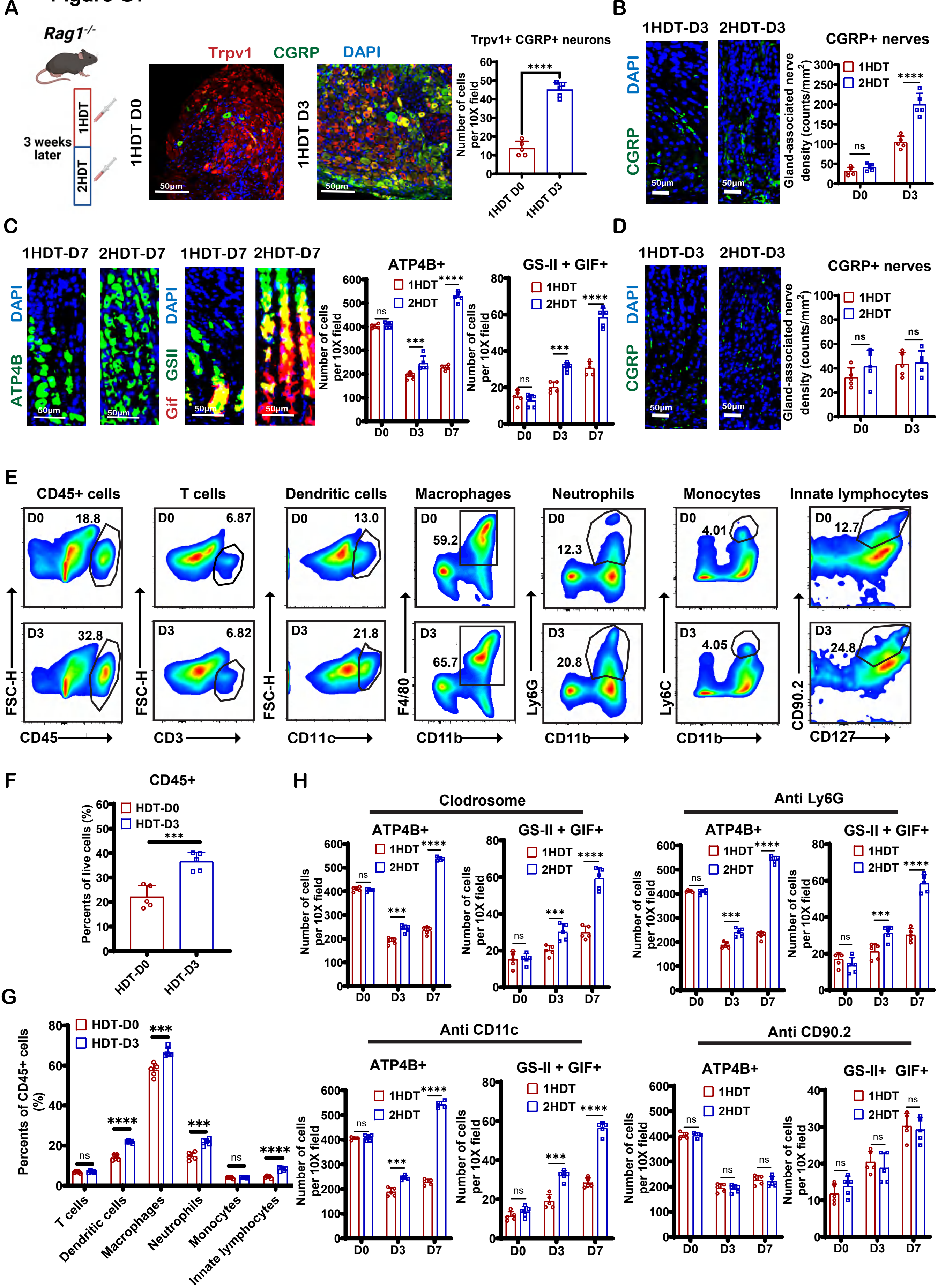

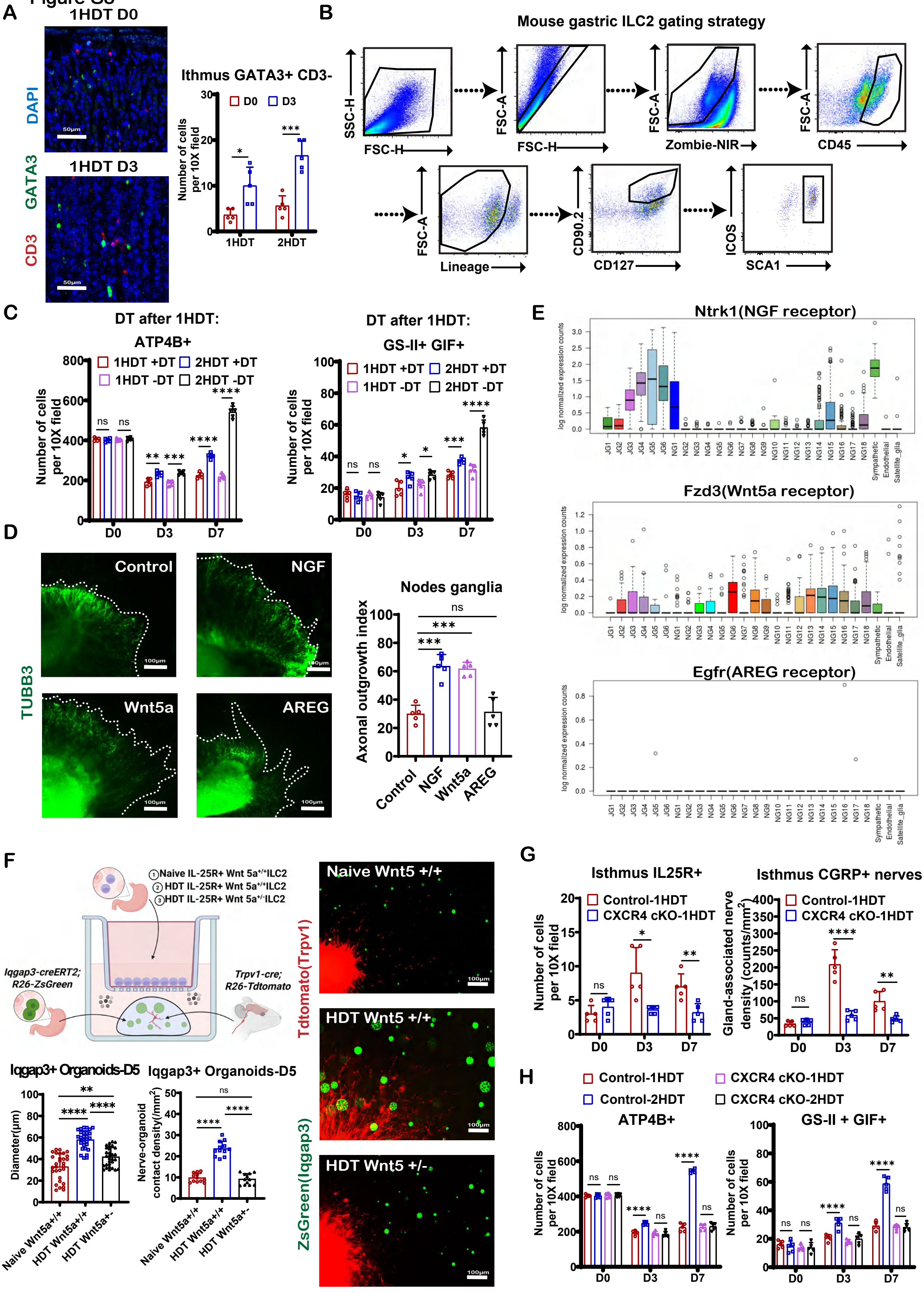

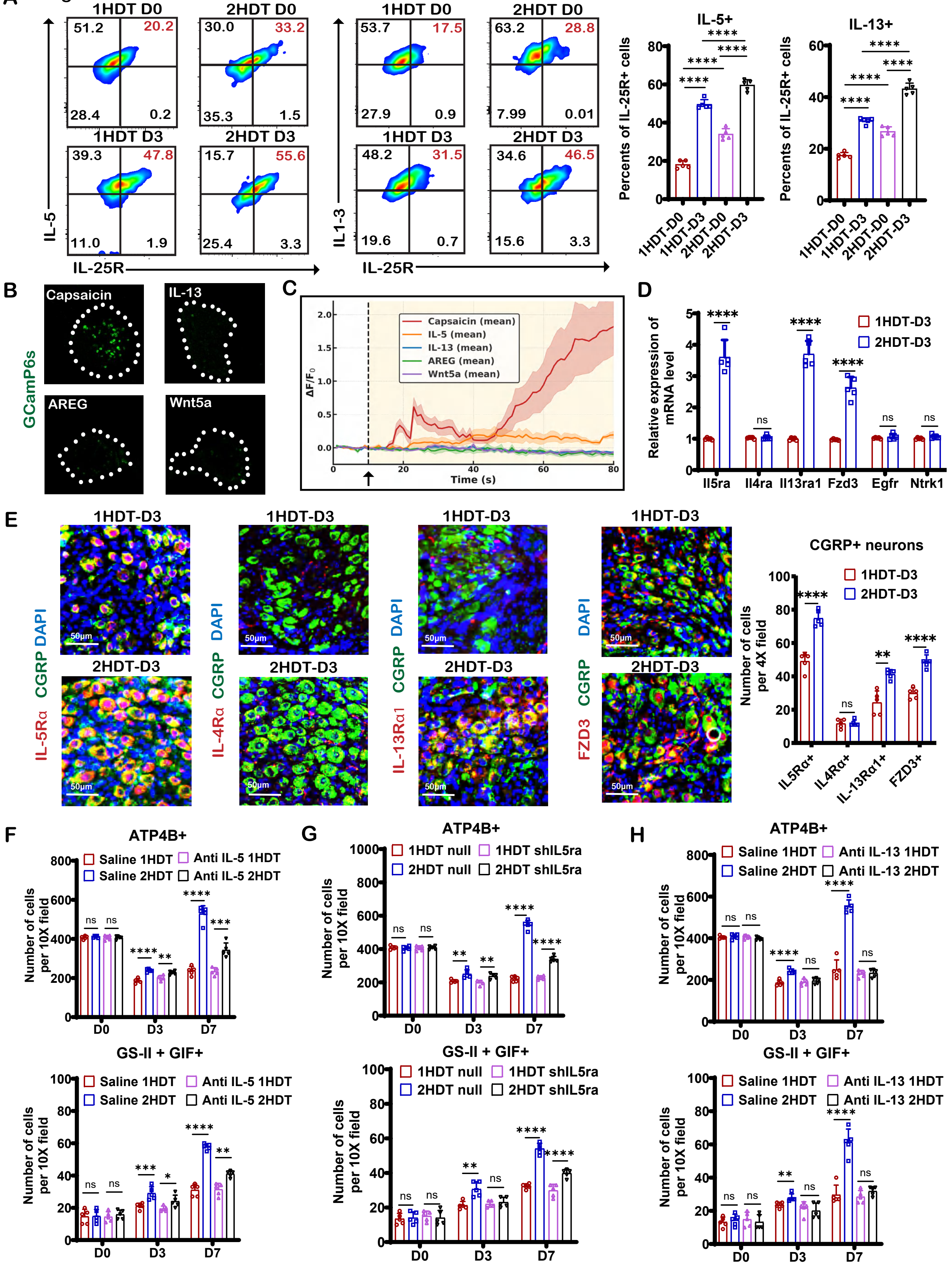

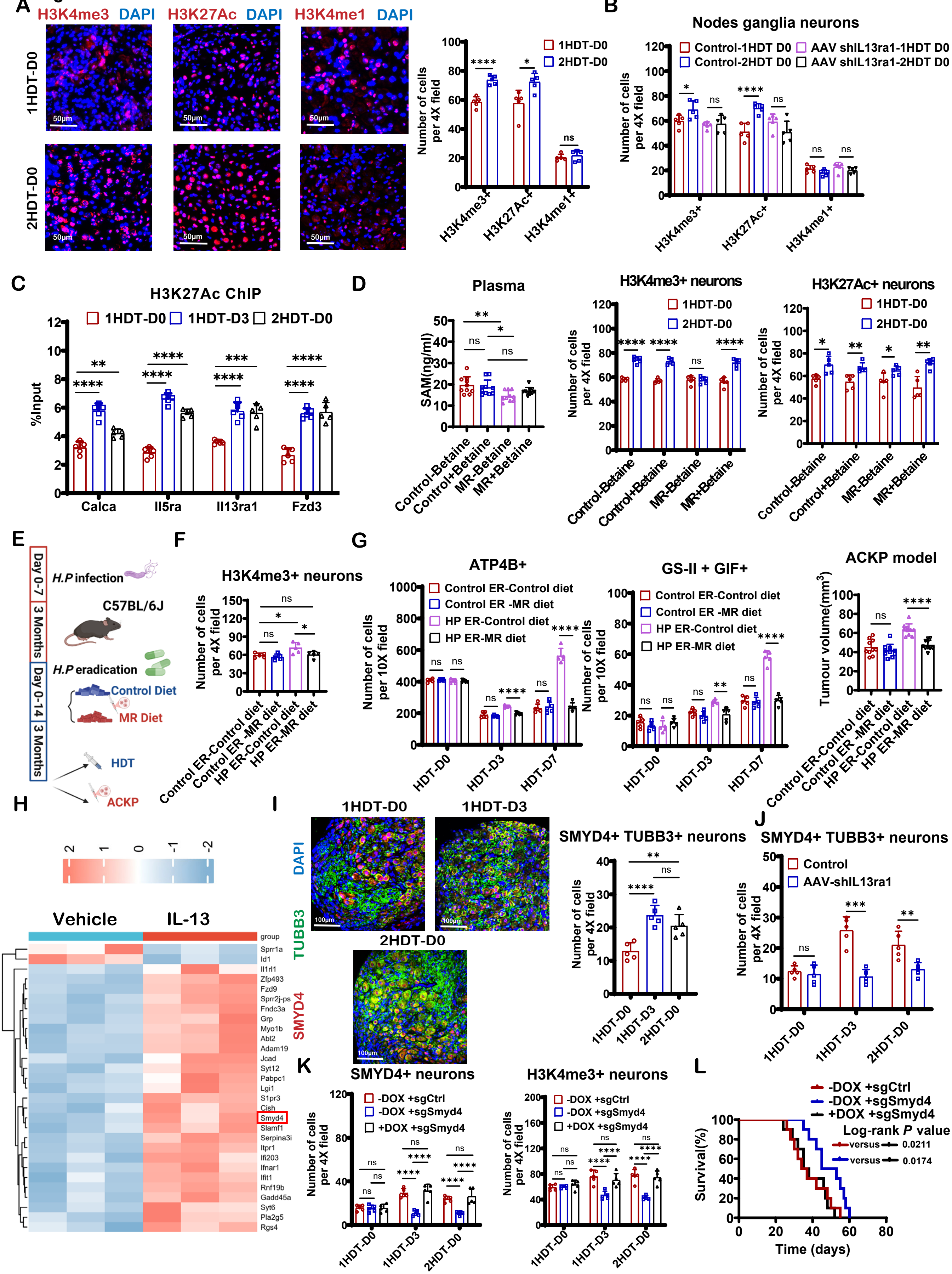
